# Cysteine oxidation triggers amyloid fibril formation of the tumor suppressor p16^INK4A^

**DOI:** 10.1101/509109

**Authors:** Christoph Göbl, Vanessa K Morris, Loes van Dam, Marieke Visscher, Paulien E. Polderman, Christoph Hartlmüller, Hesther de Ruiter, Manuel Hora, Laura Liesinger, Ruth Birner-Gruenberger, Harmjan R. Vos, Bernd Reif, Tobias Madl, Tobias B. Dansen

## Abstract

Accumulation of the CDK4/6 inhibitor p16^INK4A^ in response to oncogenic transformation leads to cell cycle arrest and senescence and is therefore frequently lost in cancer. p16^INK4A^ is also known to accumulate under conditions of oxidative stress and thus could potentially be regulated by the reversible oxidation of cysteines (redox signaling). Indeed, oxidation of the single cysteine in p16^INK4A^ in human cells occurs under relatively mild oxidizing conditions and leads to disulfide-dependent dimerization. p16^INK4A^ is an all alpha-helical protein, but here we report that upon cysteine-dependent dimerization, p16^INK4A^ undergoes a dramatic structural rearrangement and forms aggregates that have the typical features of amyloid fibrils, including binding of diagnostic dyes, presence of cross-β sheet structure, and typical dimensions found in electron microscopy. p16^INK4A^ amyloid formation abolishes its function as a CDK4/6 inhibitor. Collectively, these observations mechanistically link the cellular redox state to the inactivation of p16^INK4A^ through the formation of amyloid fibrils.

## Introduction

The *CDKN2A* gene-product p16^INK4A^ is an important cell-cycle regulator and acts as a tumor suppressor. It inhibits the D-type cyclin-dependent kinases CDK4 and CDK6 and hence prevents the downstream phosphorylation of the retinoblastoma (Rb) pocket protein ^1^. This prevents release of E2 promoter binding factor 1 (E2F1), which is otherwise required for the transcriptional regulation of proteins that regulate entry into S-phase of the cell cycle ^2^. Accumulation of p16^INK4A^ is observed upon exposure of cells to several stressors such as oxidative stress and is one of the earliest markers of oncogenic transformation ^3^. The loss of p16^INK4A^ function, or loss of Rb downstream of CDK4/6, are some of the most frequently observed mutations in tumors ^4^. Additionally, p16^INK4A^ plays an important role in aging, as clearance of p16^INK4A^-expressing senescent cells has been shown to prolong lifespan in mice ^5, 6^. The molecular basis of p16^INK4A^-mediated CDK4/6 inhibition is well established. p16^INK4A^ is a small, globular all-α-helical protein, that tightly binds into one side of the catalytic cleft of the CDK4/6 kinases. It efficiently distorts the cyclin D binding site, blocking formation of the active CDK4/6-cyclin D complex, therefore preventing Rb phosphorylation ^7, 8^. Of particular relevance for this study, the single cysteine residue (C72) present in p16^INK4A^ is located on the surface of a rigid loop and points away from the CDK4/6 kinase in the bound state; the residue is fully solvent accessible.

Reversible cysteine oxidation is the lynchpin in redox signaling, a form of signal transduction that is regulated by the cellular redox state. A more oxidizing cellular redox state, either due to elevated reactive oxygen species or a lack of reducing power, leads to the oxidative modification of specific cysteine-thiols to form reversible disulfide (S-S) bridges. These oxidative modifications can lead to structural rearrangement and can both negatively and positively regulate protein function (for a review see ^9^.

A number of observations spurred us to hypothesize that oxidation of p16^INK4A^ C72 could play a role in regulation of p16^INK4A^ activity at the molecular level. Firstly, several studies have implicated a role for increased ROS in the oncogene-induced accumulation of p16^INK4A 10^, but cysteine oxidation as the underlying mechanism has thus far not been considered nor excluded. Secondly, we identified p16^INK4A^ as prone to cysteine oxidation in a large mass-spectrometry based screen for redox sensitive proteins ^11^. Here, we provide evidence that p16^INK4A^ itself is indeed sensitive to cysteine oxidation. We found that p16^INK4A^ is readily oxidized both *in vitro* and in cultured human cells to form a disulfide-dependent homodimer, and the oxidizing conditions required are well within the physiological range. Surprisingly, this disulfide-dependent dimerization subsequently leads to the rapid formation of reversible β-amyloid fibril structures, a state that has not been previously described for p16^INK4A^. This transition leads to a loss of CDK4/6 inhibitory capacity, and this is also critically dependent on the only cysteine in p16^INK4A^. Furthermore, redox signaling-induced reversible disulfides have not been shown to induce β-amyloid fibrils before. We show that p16^INK4A^ is temporarily inactivated in response to perturbations of the cellular redox state and which has potential implications in the context of tumor biology.

Together our findings show that upon exposure to mildly oxidizing conditions the tumor suppressor p16^INK4A^ is easily converted into a functionally alternative state, characterized by an amyloid structure and disulfide-linkage. This amyloid state of p16^INK4A^ has impaired inhibitory function towards CDK4/6 activity.

## Results

### p16^INK4A^ is oxidized to form disulfide dependent homodimers in human cells

To test whether p16^INK4A^ forms disulfide-dependent complexes under oxidizing conditions, immunoprecipitation followed by non-reducing SDS-PAGE was performed. Oxidation was induced using a concentration series of hydrogen peroxide or the thiol-specific oxidant diamide (tetramethylazodicarboxamide) that ranged from subtoxic (5 µM) to mildly toxic (250 µM) in HEK293T cells. FLAG-p16^INK4A^ readily formed intermolecular disulfide-dependent complexes (Figure 1A), as judged by the large mobility shift under non-reducing conditions, that was abolished when the samples were reduced prior to SDS-PAGE (last lane). Note that only the FLAG-p16^INK4A^ disulfide-dependent dimer band is observed and that its intensity increases upon exposure to increasing oxidant concentration. If p16^INK4A^ were to undergo random crosslinking to proteins, a “smear” of disulfide-dependent complexes would be expected. We then created p16^INK4A^ mutants in which the only cysteine was replaced by alanine (C72A) or serine (C72S). We opted to use p16^INK4A^-C72A for our further experiments in human cells in order to circumvent the potential effects of the introduction of a novel phosphorylation site in the C72S mutant. The the C72A mutant is still functional as a cell-cycle inhibitor and hence the mutation does not grossly affect protein function per se (Figure S1). To further explore whether the observed intermolecular disulfide-dependent complexes are FLAG-p16^INK4A^ homodimers, Co-IP experiments were performed using combinations of wild-type (WT) p16^INK4A^ and C72A with short (FLAG) and long (mCherry) N-terminal tags. Under non-reducing conditions, bands could be observed corresponding to disulfide-dependent dimers of FLAG-p16^INK4A^-S-S-FLAG-p16^INK4A^ and of mCherry-p16^INK4A^-S-S-FLAG-p16^INK4A^ (Figure S2A) after FLAG pull down. Furthermore, diagonal electrophoresis showed that after reduction (2^nd^ dimension) FLAG-p16^INK4A^ drops out of the diagonal as a single dot, whereas a heterodimer would have revealed a second Simply Blue stained protein with similar intensity dropping out of the diagonal (Figure S2B). To test the dynamics of the intermolecular disulfide-dependent dimerization of p16^INK4A^, a time course of oxidant treatment was performed, using p16^INK4A^-C72A as a control. The oxidation of p16^INK4A^ was rapid, peaked after 10 minutes, was fully dependent on the only cysteine and was resolved upon boiling in reducing sample buffer (Figure 1B).

**Figure 1.**
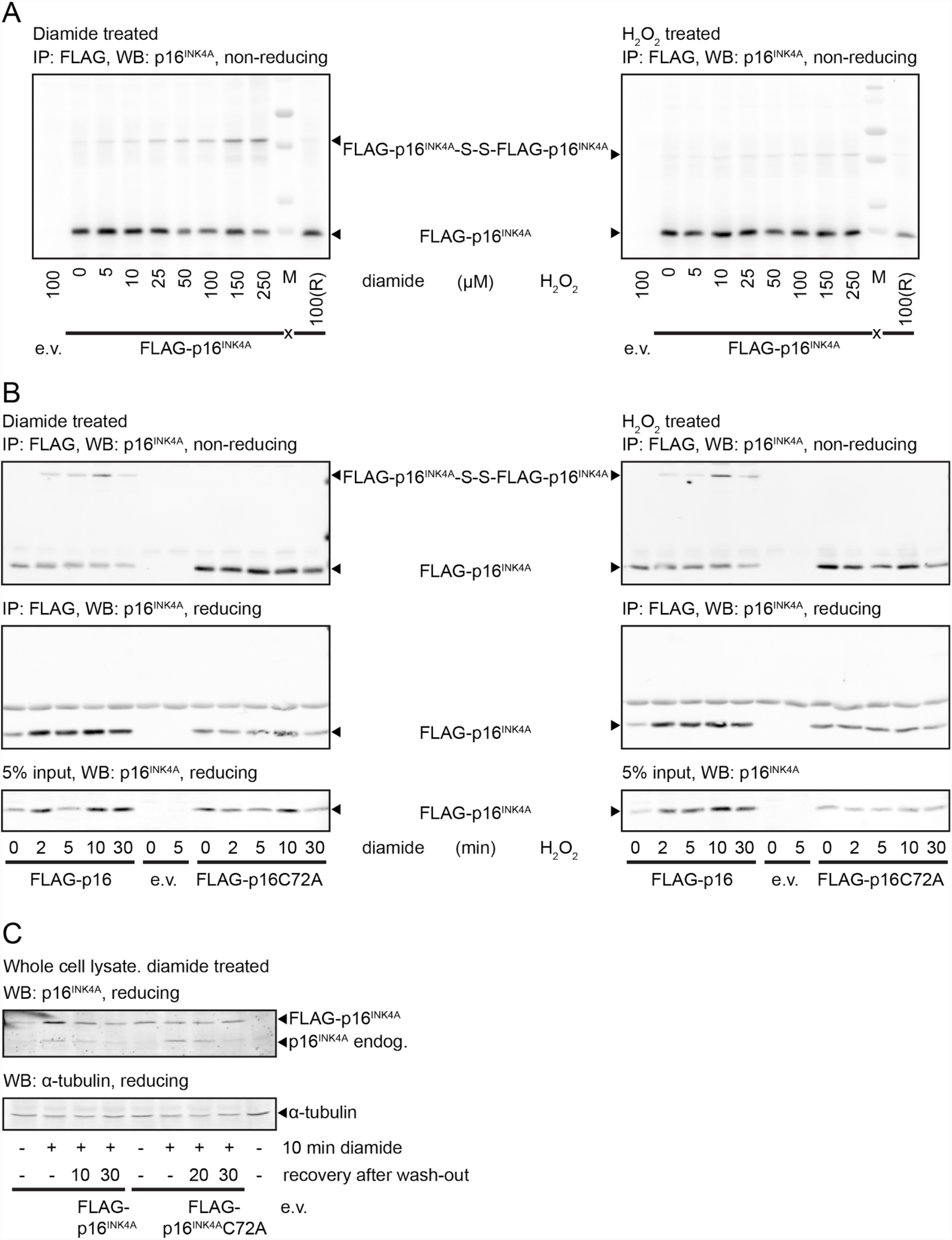
p16^INK4A^ Forms Intermolecular Disulfides Upon Exposure to Oxidants. (A) Analysis of Immuno-precipitated FLAG-p16^INK4A^ by non-reducing SDS-PAGE and Western Blot shows that part of the FLAG-p16^INK4A^ migrates at about double the molecular weight upon 5 minutes treatment with low amounts of the thiol-specific oxidant diamide (left panel) or H_2_O_2_ (right panel). Reduction (R) prior to SDS-PAGE abolishes the shift in molecular weight, indicating that it is indeed due to an intermolecular disulfide (see also Figure S2A, S2B for confirmation that the high molecular weight form of p16^INK4A^ is indeed an S-S-dependent homodimer). (B) S-S-dependent p16^INK4A^ homodimerization upon 200 µM diamide (left) or 200 µM H_2_O_2_ (right) occurs rapidly, coincides with accumulation of p16^INK4A^ protein levels and both are fully dependent on C72. (C) Endogenous p16^INK4A^ and over-expressed FLAG-p16^INK4A^ accumulate in response to 200 µM diamide with similar kinetics. Note that p16^INK4A^C72A does not accumulate whereas endogenous p16^INK4A^ does, suggesting that endogenous p16^INK4A^ levels are also regulated by Cys-oxidation. (IP: immunoprecipitation, WB: Western Blot).

Concomitantly with its oxidation, p16^INK4A^ protein abundance increased, and this was again dependent on its single cysteine residue. Protein levels already increased after a few minutes of diamide treatment, excluding a role for gene transcription but suggestive of a role for decreased protein breakdown. The accumulation is best observed using reducing SDS-PAGE (Figure 1B and 1C). The increased p16^INK4A^ protein levels rapidly returned to basal upon diamide wash-out with fresh media (Figure 1C). The experiment in Figure 1C was blotted for p16^INK4A^ rather than for FLAG, and this reveals that the endogenous p16^INK4A^ levels follow the same trend as FLAG-p16^INK4A^, whereas FLAG-p16^INK4A^-C72A levels remain unchanged, indicating that the endogenous p16^INK4A^ is also regulated through cysteine oxidation. Taken together, both the increase in p16^INK4A^ levels as well as the oxidation are dependent on C72.

### C72 of p16^INK4A^ is readily oxidized *in vitro*

To study the reactivity of the p16^INK4A^ cysteine residue directly, we expressed the recombinant protein and confirmed its correct folding by solution Nuclear Magnetic Resonance (NMR) spectroscopy. Upon treatment of the purified protein with oxidized glutathione (GSSG), we observed chemical shift changes that occurred most strongly in proximity to the cysteine residue of p16^INK4A^ (Figure 2A and 2B, Figure S3), and were reversed by addition of the reducing agent DTT, confirming the involvement of reversible cysteine oxidation. Mass spectrometric analysis of the intact protein showed that p16^INK4A^ was S-glutathionylated (S-GSHylated) after treatment with GSSG (Figure S4). S-GSHylation of the protein does not change its monomeric state or overall fold, as indicated by identical peak line shapes and unchanged elution times in size exclusion chromatography. By titrating the reduced p16^INK4A^ sample with increasing GSSG we determined the redox potential of C72 using the Nernst equation ^12^ to be −198.3 +/−1.7 meV (Figure 2C).

**Figure 2.**
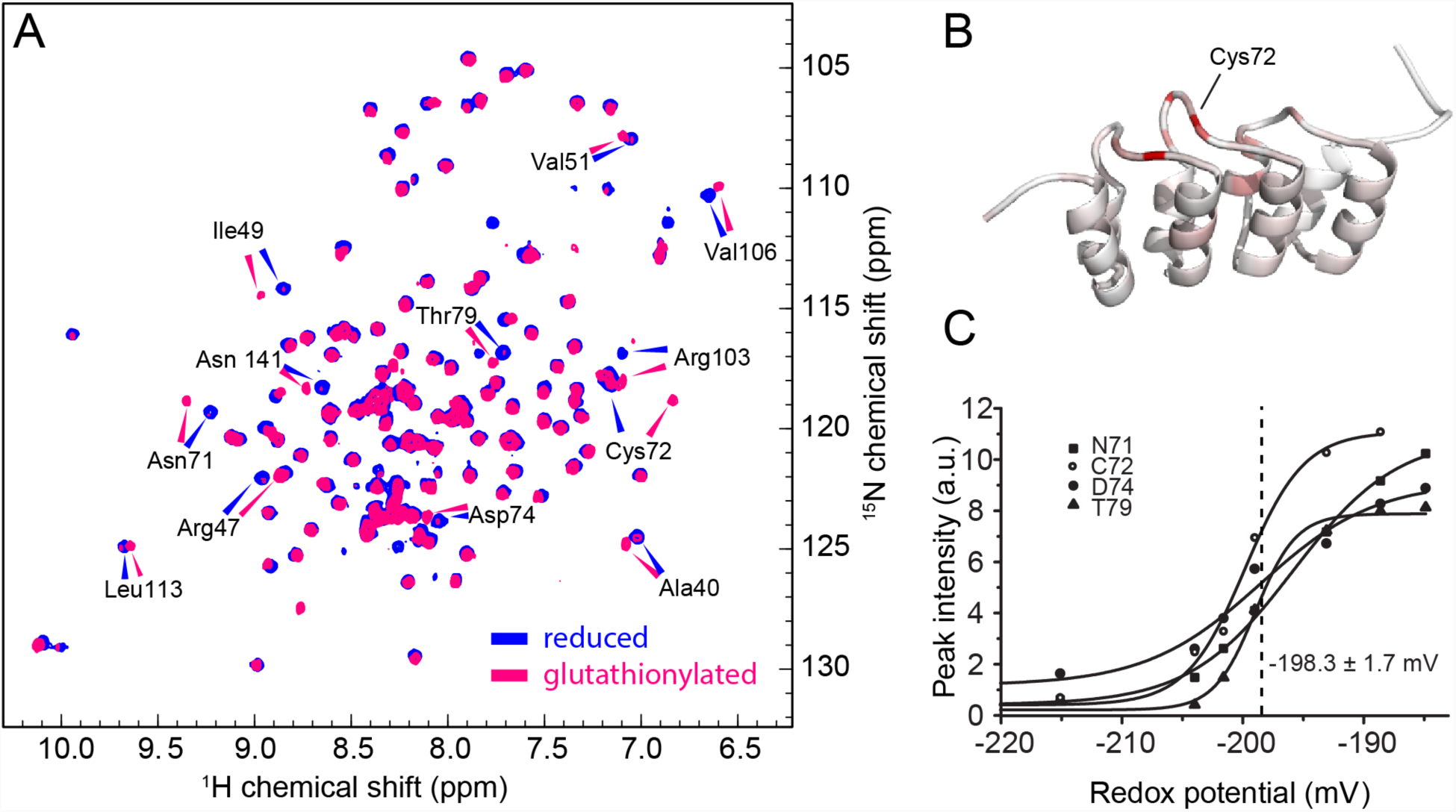
In Vitro Oxidation of p16^INK4A^. (A) ^1^H^15^N HSQC solution NMR spectrum of recombinant p16^INK4A^ in the reduced state (blue) and after S-glutathionylation (magenta). Assigned amino acids with large chemical shift changes are labeled. (B) Cartoon representation of p16^INK4A^ structure. A color gradient from white (unaffected) to red (strongly affected) shows the influence of S-glutathionylation on the chemical shift. (C) The redox potential of C72 is 198.3 ± 1.7 mV, as determined from intensity changes of four amino acids by titration of the reduced protein with oxidized glutathione.

To study the role of the cysteine residue, we used the p16^INK4A^-C72S mutant, which is the most structurally conservative mutation, differing only by replacement of the sulfhydryl group with a hydroxyl group. This mutant showed monomeric behavior during purification and largely identical chemical shifts as compared to the wild type protein (Figure S5), indicating that the mutation does not affect p16^INK4A^ protein structure per se. Addition of GSSG to this mutant did not show any indications of interaction or covalent modification. Collectively, these experiments indicate that the p16^INK4A^ C72 can be directly oxidized *in vitro* at physiologically occurring redox states.

### Disulfide-dependent homo-dimerization of p16^INK4A^ triggers aggregation

The use of GSSG as an oxidant resulted in p16^INK4A^ S-GSHylation *in vitro*, whereas experiments in human cells showed homodimerization (Figure 1) upon treatment with diamide or H_2_O_2_, even though both these agents lead to increased GSSG in cells. We therefore assessed whether H_2_O_2_ and diamide can also oxidize p16^INK4A^ *in vitro*. SDS-PAGE analysis revealed that disulfide-linked protein homodimers are also formed *in vitro*, similar to the pattern observed in the experiments in human cells (Figure S6A). In contrast to oxidation by GSSG, the addition of either H_2_O_2_ (Figure 3A) or diamide (Figure S6C and S6D) led to strong NMR chemical shift changes that indicated protein unfolding and aggregation. Comparison of size exclusion chromatography traces of reduced and oxidized samples show that oxidation of p16^INK4A^ causes a shift to high molecular weight, with the oxidized sample eluting in the void volume of the column (Figure S6B). This indicates the formation of a larger species and the absence of monomeric p16^INK4A^. Small-angle X-ray scattering (SAXS) on the oxidized samples indicated that the aggregates are larger than the detection limit of about 400 nm diameter (Figure S7).

**Figure 3.**
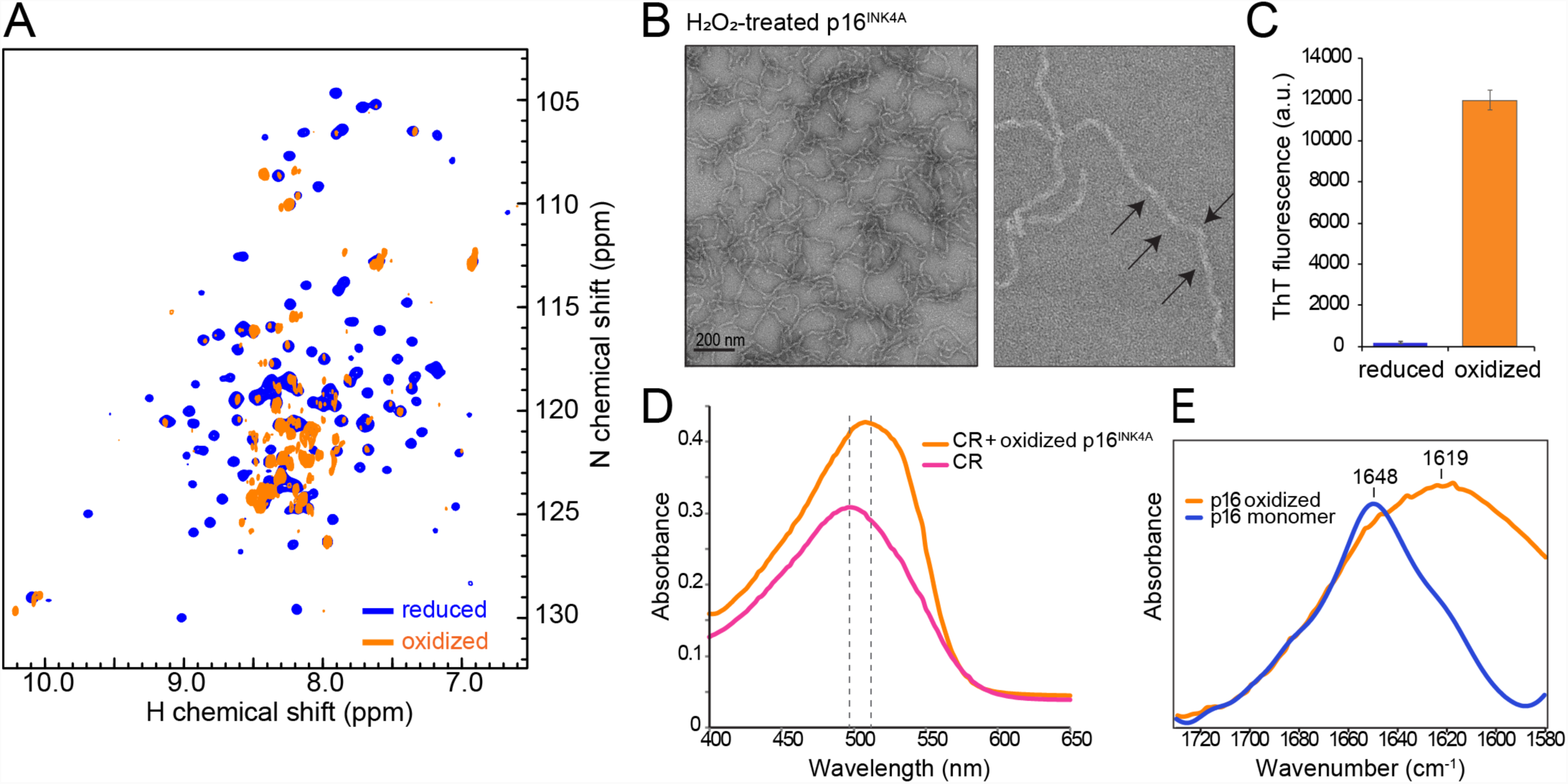
p16^INK4A^ Forms Amyloid Fibrils Under Oxidative Conditions. (A) ^1^H,^15^N HSQC solution NMR spectra of recombinant p16^INK4A^ before (blue) and after oxidation with 50 mM H_2_O_2_ for 10 h at room temperature (orange). The collapse of the peaks to the center of the spectrum suggests formation of unstructured or aggregated protein. (B) Negative-stained transmission electron micrographs of diamide-treated p16^INK4A^ showing the presence of amyloid fibrils. Right panel is at higher magnification, arrows highlight the twisted morphology which is typical of amyloid fibrils. (C) Fibril formation of p16^INK4A^ monitored by thioflavin-T fluorescence measurement. Error bars represent standard deviation from four measurements. (D) Congo red absorption maximum shifts when bound to oxidized p16^INK4A^, indicating the presence of amyloid fibrils. (E) Amide I region of the attenuated-total-reflectance Fourier-transform infrared spectra of p16^INK4A^ monomer and fibrils, with peak maxima indicating structure contain primarily alpha helix and cross-beta sheet secondary structure, respectively.

S-Glutathionylation of p16^INK4A^ C72 would abolish its reactivity towards diamide or H_2_O_2_. To exclude that the observed aggregation of p16^INK4A^ stems from modifications other than C72, we also performed NMR spectroscopy on the S-glutathionylated p16^INK4A^ (GS-p16^INK4A^) treated with diamide or H_2_O_2_. No structural changes were observed and the monomeric, S-glutathionylated protein sample was stable in presence of the same concentrations of oxidizing agents (Figure S8). Together, these experiments indicate the formation of an aggregated p16^INK4A^ species upon intermolecular disulfide bond formation *in vitro*.

### Disulfide-linked homodimers of p16^INK4A^ form β-amyloid fibrils

Based on the indications of aggregation, transmission electron microscopy was employed to study the nature of the p16^INK4A^ aggregates that formed upon oxidation. Interestingly, we found that the oxidized p16^INK4A^ samples produced fibrillar structures (Figure 3B). One prominent group of fibrillar proteins are amyloid fibrils, which are characterized by a cross-β sheet core structure, meaning that β-strands run perpendicular to the fibril long axis ^13, 14^. To determine if p16^INK4A^ fibrils are amyloid, we used the amyloid-specific dyes Thioflavin T (ThT) and Congo Red ^15-17^. ThT assays showed the characteristic increase in fluorescence when bound to oxidized p16^INK4A^ (Figure 3C), while Congo Red absorbance measurements showed the red shift and increased absorbance characteristic of amyloid binding (Figure 3D). Fourier-transform infrared spectroscopy can distinguish cross β-sheets found in amyloid fibrils from β-sheets found in globular monomeric proteins ^18, 19^; in line with the dye-binding data, p16^INK4A^ fibrils had a maximum at 1620 cm^-1^, which falls within the typical amyloid β-sheet region, confirming the amyloid structure of the fibrils (Figure 3E). We performed a computational analysis using four fibril prediction algorithms ^20-23^ and all suggested a high propensity in the region from residue no. 91-99 (Figure S9A).

### Solid-state NMR spectroscopy confirms p16^INK4A^ forms β-amyloids

Solid-state NMR spectra were recorded on a uniformly ^13^C-labelled p16^INK4A^ sample to confirm the secondary structure of the aggregated state. In ^13^C-^13^C correlation spectra, only amino acid residues within a structured core are visible (Figure 4A). We detected a well-dispersed spectrum indicating a structured region of approximately 20-30 residues. Certain residue types are absent from the spectrum, for example isoleucine (Figure S9B), confirming that only a subset of the sequence is present in the structural core of the amyloid fibrils. Many of the peaks could be assigned to their amino-acid type due to their distinctive Cα and Cβ peak positions, which allowed for determination of the secondary structure. All assigned peaks have peak positions that are characteristic for β-sheet or random coil (Figure 4B), confirming that p16^INK4A^ aggregates have a β-sheet core and are therefore amyloid. These findings emphasize the dramatic structural rearrangement from the monomeric eight-alpha-helical bundle.

**Figure 4.**
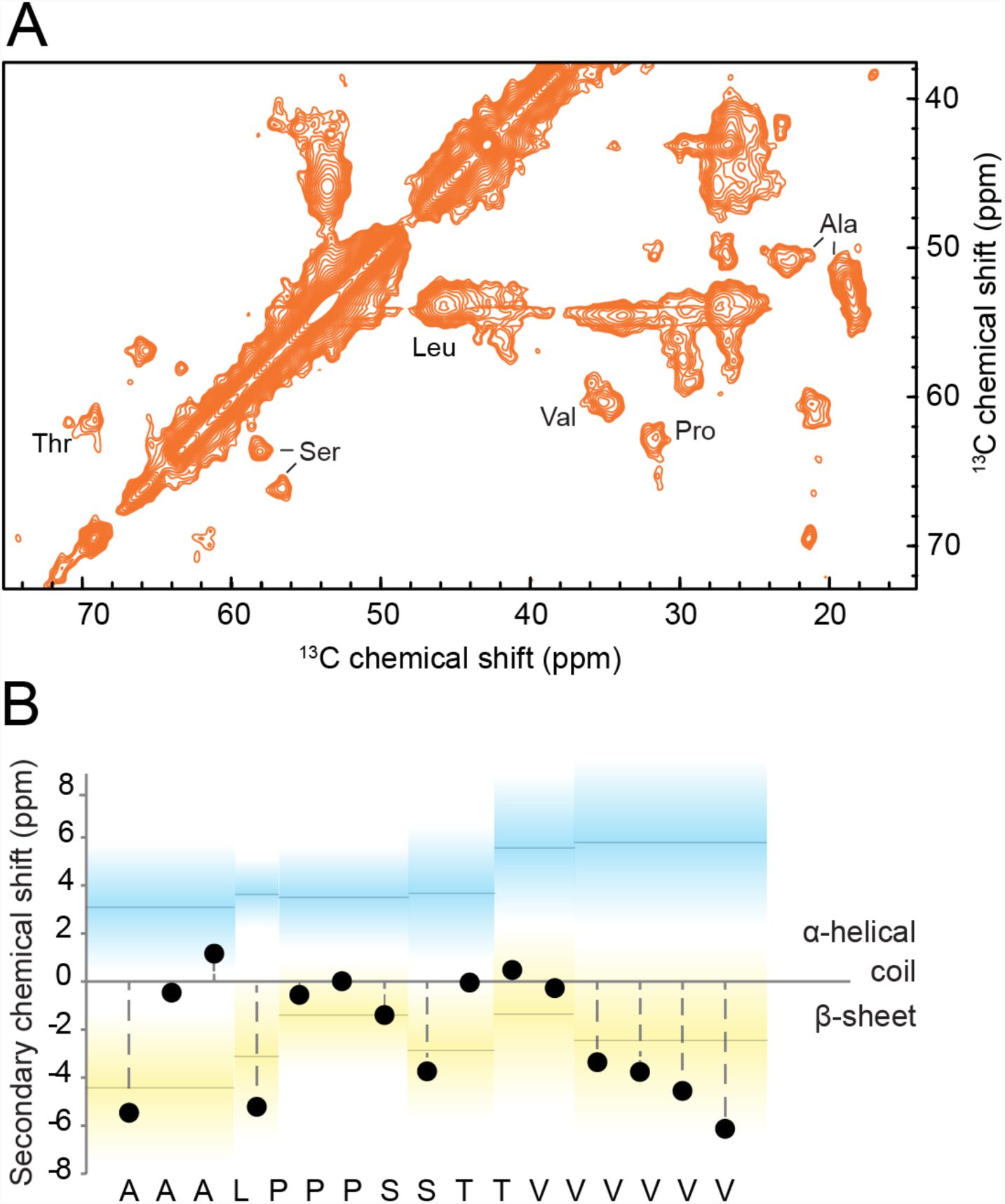
Solid State NMR Spectroscopy Analysis of Amyloid p16^INK4A^. (A) ^13^C-^13^C correlation magic-angle-spinning solid state NMR spectrum of p16^INK4A^ amyloid fibrils. Assigned amino acid types are indicated. The spectrum was recorded using a proton-driven spin diffusion (PDSD) pulse sequence with 50 ms mixing time. (B) Secondary chemical shift of uniquely assigned amino acid types showing the presence of β-sheet secondary structural elements.

### Intermolecular p16^INK4A^ disulfide bond formation is essential for amyloid fibril assembly

We further investigated the role of oxidation of the cysteine residue in fibril formation by comparing the WT protein to p16^INK4A^-C72S. Electron microscopy revealed that upon treatment with as little as 200 µM of H_2_O_2_ or diamide, WT samples formed large fibrillar structures (Figure 5A, top). There was no discernable difference between the morphology of fibrils formed with diamide or hydrogen peroxide treatment, or when higher concentrations of oxidizing agent were used. p16^INK4A^-C72S samples displayed a very different morphology, instead forming unordered, amorphous aggregates, as well as a small amount of short fibrillar species (Figure 5A, middle). Glutathionylated-p16^INK4A^ (GS-p16^INK4A^) samples were also analyzed by electron microscopy, and were found to form large, granular aggregates without any fibrils upon treatment with either H_2_O_2_ or diamide (Figure 5A, bottom). Although aggregates are seen by electron microscopy, the NMR spectra of GS-p16^INK4A^ indicate the sample is largely soluble and monomeric (Figure S7), suggesting that the aggregates seen by electron microscopy constitute only a small proportion of the total protein.

**Figure 5.**
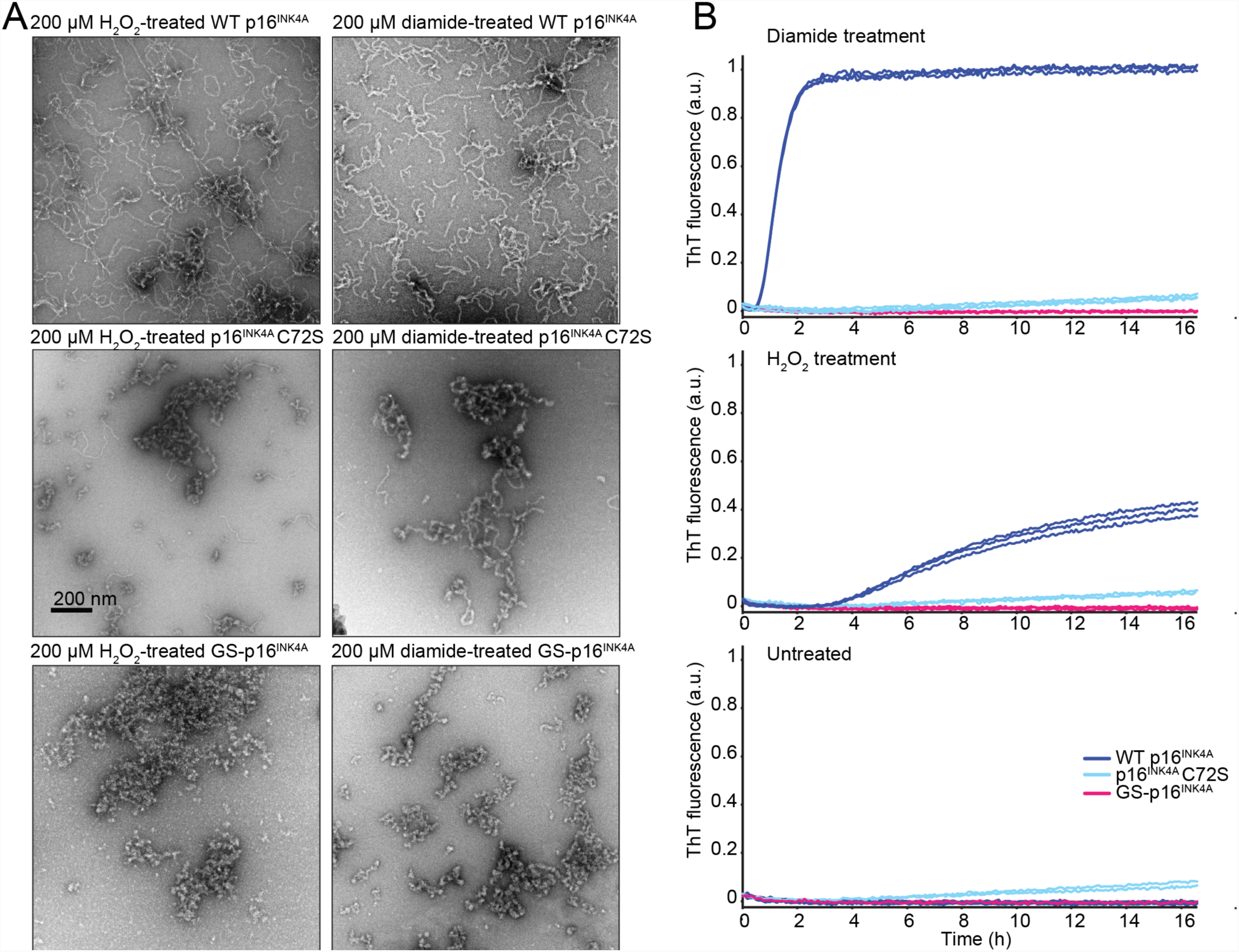
C72 Is Important For Fibrillar Morphology And ThT Binding. (A) Negative-stain transmission electron micrographs of hydrogen peroxide (left) and diamide (right) treated p16^INK4A^. Top: WT p16^INK4A^. Middle: p16^INK4A^C72S. Bottom: GS-p16^INKA^. (B) Thioflavin-T fluorescence kinetics assay of p16^INK4A^ samples with diamide addition (top) H_2_O_2_ addition (middle) and without oxidizing agents (bottom). The fluorescence signal of samples of p16^INK4A^ (WT, C72S and GS), supplemented with Thioflavin-T, was monitored over time and in the presence or absence of diamide and H_2_O_2_. Three replicates were measured per sample and each are separately plotted.

We next investigated the influence of intermolecular disulfide bond formation on the kinetics of fibril formation, by measuring the time- and oxidation-dependence of WT, C72S- and GS-p16^INK4A^ variants using the ThT assay (Figure 5B). Upon treatment of WT p16^INK4A^ with oxidizing agents, the sample fluorescence rapidly increased as ThT-positive fibrils were formed, with maximum signal being achieved within 2 hours upon 200 µM diamide treatment (Figure 5B, top) and more than 16 hours for 200 µM H_2_O_2_ treatment (Figure 5B, middle), while no fibrils were formed in the absence of oxidizing agents (Figure 5B, bottom). The C72S mutant displayed only a very slight increase in fluorescence over time, and this was not affected by the presence or absence of oxidizing agents. GS-p16^INK4A^ did not form ThT-positive aggregates under any condition. These results emphasize that the amorphous aggregates observed by EM for the p16^INK4A^-C72S and GS-p16^INK4A^ are not amyloid, both by morphology and by dye-binding properties. Therefore, we conclude that the intermolecular disulfide bond formation of p16^INK4A^ through C72 is critical for the formation of p16INKA amyloid fibrils.

### p16^INK4A^ cysteine oxidation triggers aggregation in cultured cells

Having shown that p16^INK4A^ forms amyloid fibrils upon oxidation *in vitro*, we investigated to what extent this behavior was conserved in cultured human cells. HEK293T cells expressing FLAG-tagged p16^INK4A^ WT and C72A mutant were exposed to diamide and processed for a filter trap assay: an assay that is commonly used to detect amyloids in cell extracts (Scheme top of Figure 6A). Exposure to diamide prior to lysis rapidly led to trapping of p16^INK4A^ on the filter membrane, and this was dependent on C72. Reduction of the protein lysates with DTT prior to washing abolished trapping, suggesting that the aggregation was reversible. The loss of p16^INK4A^ aggregates was also apparent upon recovery from the exposure to diamide: after 60 minutes the aggregates were largely resolved (Figure 6A), reminiscent of turnover of the disulfide-linked homodimer in Figure 1C. Of note, the amyloids formed *in vitro* were also reversible: boiling in non-reducing sample buffer for five minutes prior to SDS-PAGE and Western blot resulted mainly in detection of the S-S-dependent homodimeric p16^INK4A^ (Figure S10).

**Figure 6.**
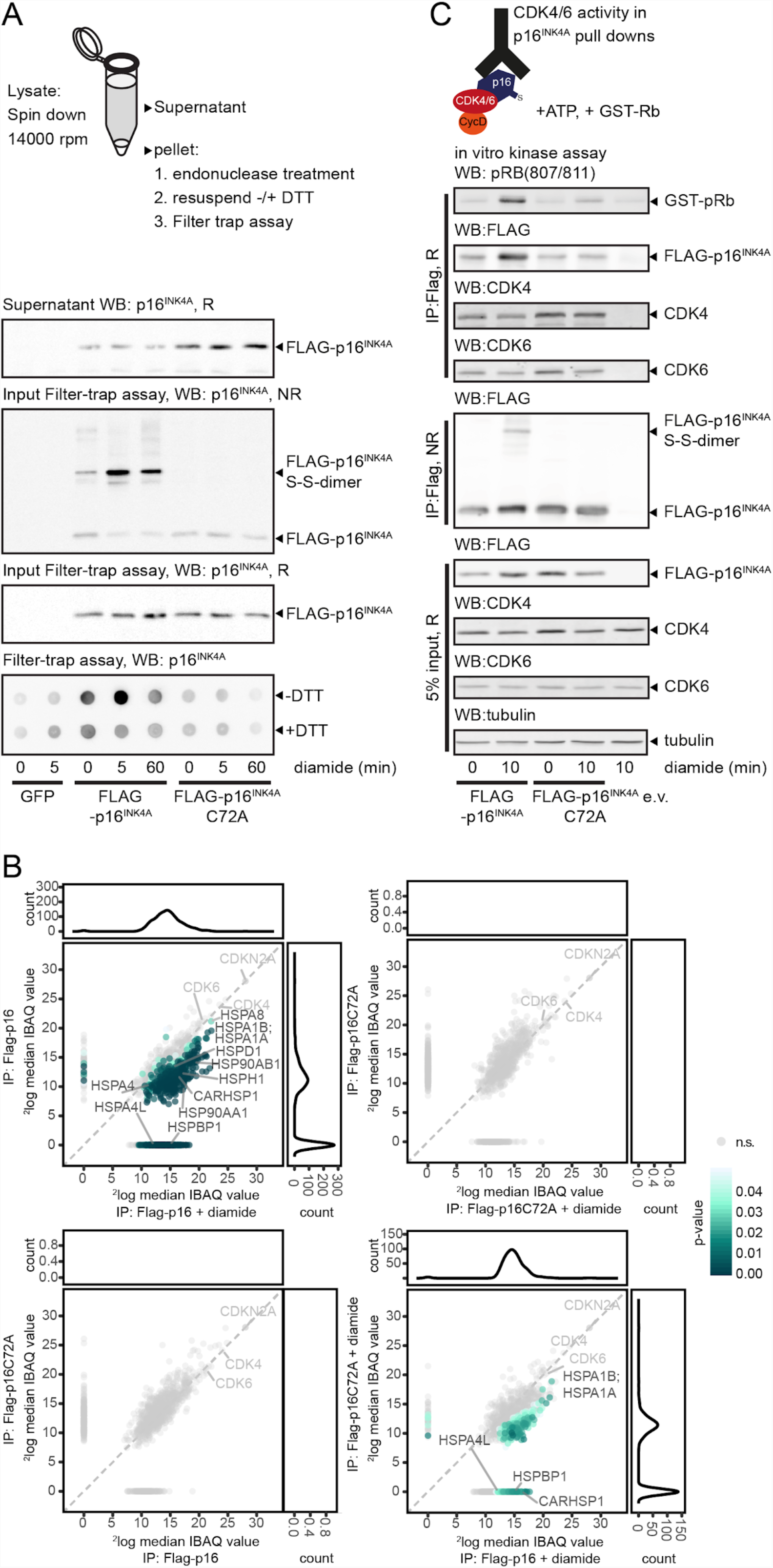
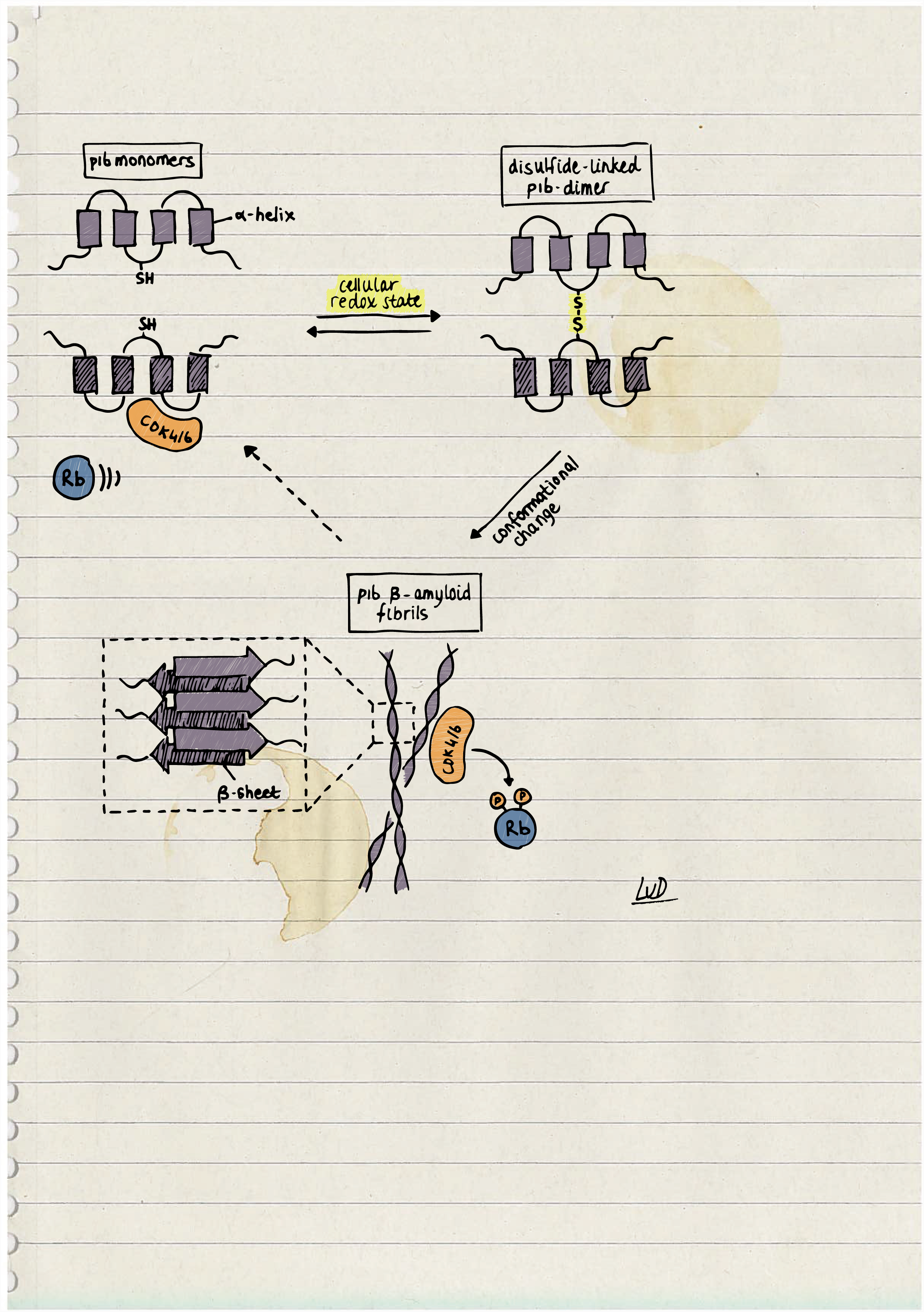
p16^INK4A^ Aggregates in Live Cells in Response to Oxidation. (A) Filter trap assay for the detection of aggregates in cell lysates. Note that the majority of p16^INK4A^ in the pellet is in the form of-S-S homodimers after lysis. p16^INK4A^, but not p16^INK4A^C72A, was trapped on the filter membrane upon treatment with 200 µM diamide and trapping was prevented by pretreatment with DTT. Equal amounts of p16^INK4A^ and p16^INK4A^C72A were used as input for the filter trap assay. (B) MS/MS analysis of the p16^INK4A^ interactome upon oxidation. Diamide (200 µM) treatment induces large changes in proteins that interact with p16^INK4A^ and this largely depends on C72. Note that the interaction with CDK4 and CDK6 is not altered by diamide treatment or C72 dependent and that equal amounts of p16^INK4A^ and p16^INK4A^C72A were pulled down. Several chaperone proteins can be found to interact in an oxidation dependent manner. Log-transformed median IBAQ values are plotted, colors represent proteins with adjusted p-values smaller than 0.05. Marginal plots represent the total number of significant proteins. n=6 biological replicates for p16^INK4A^ and p16^INK4A^C72A, n=5 for p16^INK4A^+diamide and n=4 for p16^INK4A^C72A +diamide (C) In vitro CDK4/6 kinase assay on p16^INK4A^ pulldowns shows that oxidation of p16^INK4A^C72 impairs its inhibitory function. Note that oxidation or mutation of p16^INK4A^C72 does not affect the amount CDK4/6 that is co-immunoprecipitated. (R: Reducing, NR: Non-reducing)

The phase-transition from monomeric to the amyloid form of p16^INK4A^ would likely change its biochemical properties and protein-protein interactions. To test this, the interactomes of immunoprecipitated FLAG-p16^INK4A^ and FLAG-p16^INK4A^C72A were investigated and compared by label-free quantitative mass spectrometry. Under basal conditions the interactomes were largely identical, but this changed dramatically upon diamide treatment (Figure 6B). Diamide-treated WT p16^INK4A^ showed a large number of binding partners not identified in diamide-treated p16^INK4A^-C72A pull-downs. The interaction with CDK4 and CDK6 was not affected by diamide treatment or the C72A mutation. Proteins that form amyloids are known to bind to chaperones and indeed several Heat Shock Proteins were enriched in WT p16^INK4A^ pull downs upon diamide treatment, again indicative of aggregate formation (Figure S11).

### p16^INK4A^ cysteine oxidation restores CDK4/6 kinase activity

We then investigated how the p16^INK4A^ function as a CDK4/6 inhibitor would be affected by amyloid formation. CDK4/6 was co-immunoprecipitated with FLAG-tagged p16^INK4A^, and *in vitro* kinase assays were performed using a GST-tagged Rb fragment as a substrate (Scheme Figure 6C). These experiments revealed that oxidation of WT p16^INK4A^ or p16^INK4A^-C72S did not greatly affect the binding to CDK4/6 (Figure 6C), which agrees with our observations in the MS experiment (Figure 6B). The in vitro kinase assay showed that p16^INK4A^-associated CDK4/6 was, as expected, inactive under basal conditions. CDK4/6 activity towards the GST-Rb fragment was regained when cells were exposed to diamide prior to lysis, despite the increased amount of p16^INK4A^ that was pulled down. The reactivation was strictly dependent on p16^INK4A^ C72. Taken together, these observations suggest that the amyloid form of p16^INK4A^ still interacts with CDK4/6 but in a manner that does not lead to kinase inhibition.

## Discussion

Amyloid fibrils are structures that can be formed by a wide variety of protein sequences ^24^. These fibrils share common features, including an unbranched fibrillar morphology and a cross-β sheet core structure. Amyloids were previously thought to be largely disease-related, especially involved in neurodegenerative diseases, but there are increasing cases of amyloid fibrils discovered that have a biological function. Several mammalian examples have been reported in recent years, including the melanosome protein pMeL, amyloids involved in hormone storage, and the RIPK proteins, whose fibril formation appears to trigger necroptotic signaling pathways ^25-28^ In this work we show that p16^INK4A^ can form fibrils under physiological conditions and in cell-based models, and that the formation is triggered by oxidation of the single cysteine residue and homodimer formation. We presented evidence that p16^INK4A^ can form aggregates that have the typical features of amyloid fibrils, including binding of diagnostic dyes, presence of cross-β sheet structure, and typical dimensions found in EM. The critical dependence of a disulfide cross-linked dimer as a subunit has not been observed so far and highlights the role of the cellular redox state as an important regulator of fibril formation.

The role of cysteine chemistry in amyloid formation has been discussed in detail previously ^29^. Most commonly, disulfide bonds are found to stabilize a soluble, usually monomeric, form of the protein, preventing its un- or misfolding and aggregation. In many cases, blocking of disulfide bond formation by mutagenesis or chemistry generates proteins that are more prone to amyloid formation, such as has been demonstrated for lysozyme ^30^, SOD1 ^31, 32^, insulin ^33^ and prion protein ^34, 35^. In contrast, oxidation of the single cysteine of p16^INK4A^ is important for the transition of the monomeric protein to form amyloid fibrils, and modification of the cysteine by mutagenesis or S-glutathionylation prevents its amyloid formation. Proteins in which disulfide bonds have been postulated to be involved in stabilization of amyloid fibrils include the prion protein and RIPK fibrils, however disulfide bond formation itself is not required for conversion to amyloid in either of these cases ^34-36^.

Several properties of p16^INK4A^ fibrils fit with those expected for functional amyloids, including their rapid formation (disease-related amyloids can take days to form fibrils) ^37, 38^, and their lack of clear polymorphism (no evidence of peak doubling in the solid-state NMR spectrum) ^39, 40^. One striking difference in the properties of p16^INK4A^ amyloid fibrils, compared to classical disease-linked amyloids like amyloid-β and α-synuclein, is the apparent reversibility of the fibrils. Typical amyloids are highly stable and are not disaggregated by SDS buffers and do not run into SDS-PAGE gels ^24, 41, 42^. Our results show that SDS is able to disaggregate p16^INK4A^ fibrils into dimers, and that reducing-agent treatment further returns p16^INK4A^ to a monomeric state. This was shown for both recombinantly-produced p16^INK4A^ amyloid *in vitro*, as well as p16^INK4A^ amyloid formed in a cellular model. Furthermore, we find that p16^INK4A^ amyloids are transiently formed and do not persist in the cellular environment. This result potentially adds to an emerging pattern of lability of amyloid fibrils that have functional roles in the cell. For example, RIPK1/3, and the phase-separating proteins FUS and hnRNPA2 are all reported to form functional amyloid fibrils, and are apparently SDS soluble ^36, 43^. This may be a hallmark for functional fibrils, which must be regulated and removed when the function is no longer needed. Our observations would also fit with a model in which p16^INK4A^ is stored in an inactive state in the form of reversible amyloid fibrils to temporarily relieve CDK4/6 inhibition, while keeping the pool of p16^INK4A^ available for reuse rather than targeting it for degradation. In contrast, highly stable, persistent aggregates may have toxic properties, such as in the case of Tau or TDP-43 aggregates, which are pathogenically related to various dementias and amyotrophic lateral sclerosis, respectively ^44, 45^.

It is now generally accepted that the cellular redox environment contributes to various signaling pathways and that H_2_O_2_ can act as a secondary messenger molecule ^9^. Cysteine residues are frequently found to be modified in large-scale analysis of the cysteinome and thousands of proteins are affected by redox regulation ^46, 47^. Redox potentials vary widely between cell types and specific compartments, ranging from −374 mV of the strongly reducing NADPH/NADP^+^ redox couple in the liver cytosol to −180 mV from the GSH/GSSG couple in the oxidizing endoplasmic reticulum of B-cells ^48^. In addition, the average redox potential varies greatly between cell cycle stages ^49^. The redox potential of −198 mV we found for the oxidation of C72 of p16^INK4A^ is well within the physiological range. One difference between *in vitro* and cell culture oxidation of p16^INK4A^ is that *in vitro* p16^INK4A^ could be stably GSH-ylated whereas in cell cultures we only observed the S-S homodimer. This might seem surprising, because there is ample GSH available in the cell and addition of micromolar ranges of diamide will likely react to form GSSG. It therefore may be unlikely that in cells diamide and H_2_O_2_ oxidize p16^INK4A^ directly, but rather that its oxidation is the result of a redox relay or disulfide exchange mechanism, like has been proposed for other redox-sensitive proteins ^50^. However, we cannot exclude that transient GSH-ylation of p16^INK4A^ occurs in cells. Of note, tumor cells have been frequently reported to exhibit a more oxidizing milieu ^51^. This could mean that p16^INK4A^ would be more often in the dimerized and aggregated state, and that this could contribute to aberrant perturbation of the S-phase checkpoint in tumors that express p16^INK4A^. Amyloid formation-dependent inactivation of tumor suppressor proteins has been described before for p53 and also in this case amyloid formation was reversible ^52^. The observations that p16^INK4A^ levels are often increased in tumors might not only be due to transcriptional upregulation, but may in addition be the result of a slower turnover of p16^INK4A^ in the aggregated state. We observed that accumulation of p16^INK4A^ upon oxidizing conditions is too fast to depend on transcription, which may support the hypothesis that the amyloid state is causing accumulation of p16^INK4A^ due to less efficient degradation. It remains to be investigated to what extent p16^INK4A^ inactivation by amyloid formation indeed contributes to oncogenic transformation.

## Acknowledgements

We would like to thank Marc Pagès-Gallego, Vladimir Hristov, Dr. Holger Rehmann, Dr. Marten Hornsveld, Zoë Alexiou and Kim van Dorenmalen for assistance with preparation and validation of DNA constructs and cell lines, additional experiments, analysis and discussion. We thank Dr. Mark Hipp and Dr. Willianne Vonk for discussions on aggregation and help with the Filter trap assay protocol. We are grateful for input of the Burgering and Dansen lab members on this project. We would like to gratefully acknowledge the assistance of Dr. Carsten Peters and Prof. Sevil Weinkauf with electron microscopy measurements, and the assistance of the groups of Prof. Johannes Buchner and Prof. Stephan Sieber with the use of the fluorescence plate readers. We would also like to thank Dr. Riddhiman Sarkar with assistance with solid-state NMR measurements, and Dr. Ralf Stehle for assistance with FTIR measurements. This work was supported by the President’s International Fellowship Initiative of CAS (No. 2015VBB045, to TM), the National Natural Science Foundation of China (No. 31450110423, to TM, KLI425 and KLI645), the Austrian Science Fund (FWF: P28854 and I3792 to TM), the Austrian Research Promotion Agency (FFG: 864690), the Integrative Metabolism Research Center Graz, the Austrian infrastructure program 2016/2017, BioTechMed-Graz, and the OMICS center Graz. CG was supported by a PFP fellowship of the Helmholtz Zentrum München. VM was supported by a CJ Martin Early Career Fellowship from the National Health and Medical Research Council of Australia. TD and MV were supported by a grant from the Dutch Cancer Society (KWF Kankerbestrijding). HV was supported by the ProteinsAtWork research programme.

## Author Contributions

C.G., V.M., T.M. and T.D. planned the study, performed experiments and wrote the manuscript with input from the other authors. P.P., L.D., M.V., and H.R. performed immuno-precipitation and Western blot experiments in cultured cells. M.V. and P.P. did the in vitro kinase assay. The filter trap assay was done by P.P. Mass spectrometry sample preparation, measurement and analysis for the experiments using cultured cells was done by L.D., M.V. and H.V. C.G. produced recombinant protein samples, C.G., C.H. and T.M. performed solution NMR experiments, V.M. and M.H. measured electron microscopy, V.M and B.R. measured solid-state NMR experiments, and V.M. performed all other amyloid characterization experiments. L.L. and R.B.G. performed intact mass spectrometry measurements. R.B.G and B.R. edited the manuscript. L.D. sketched the graphical abstract.

## Methods

### Cell lines, plasmids, antibodies and compounds

Cells were cultured in DMEM low glucose (HEK293T) supplemented with FCS (10%), L-glutamine (2mM) and Penicillin-Streptomycin (100 Units-0.1mg per ml medium) (all from Lonza). Cells were transfected using PEI (Sigma Aldrich) or X-tremeGENE 9 (Roche) according to manufacturer’s instruction.

The vector containing the truncated form (Δ1–8) of wild type p16^INK4A^ was a gift from R. Medema ^53^. The cysteine mutant of p16^INK4A^ was created by site-directed pcr mutagenesis. mCherry-(backbone pLV-CMV-bc) and FLAG-His-tagged(backbone Pcdna3) p16^INK4A^ constructs were created using gateway technology (Life Technologies). For cell cycle profiling, wild type and C72A p16INK4A were cloned in the pTON vector, which was a gift of M. Tanenbaum ^54^.

The following antibodies were used in this study: FLAG (M2 F3165 Sigma Aldrich), p16^INK4A^ (10883 Proteintech), p16^INK4A^ (ab16123 Abcam), CDK4 (C-22 Santa Cruz), CDK6 (C-21 Santa Cruz), tubulin (CP06 Merck Millipore), Phospho-Rb Ser807/811 (9308 Cell Signaling).

Solutions/compounds not mentioned elsewhere: 35% hydrogen peroxide (Merck).

### Co-immunoprecipitation, SDS-PAGE and Western Blotting

Transfection was performed two days prior to sample preparation. Cells were scraped in 100 mM N-ethyl-maleimide in PBS (37 °C) to prevent post-lysis oxidation and collected by centrifugation at 1200 rpm for 3 minutes. Cells were lysed in a standard lysis buffer containing 50 mM Tris pH 7.5, 1% triton X-100, 1.5 mM MgCl_2_, 5 mM EDTA, 100 mM NaCl supplemented with Aprotinin, Leupeptin and NaF. 100 mM iodoacetamide was added to to prevent post-lysis oxidation of cysteines. The cell lysates were subsequently centrifuged at 14000 rpm for 10 min. 50 µl of the supernatant was kept as input and the remaining supernatant was used for immunoprecipitation with the indicated antibody coupled to protein A-Agarose (Roche) or in case of FLAG with anti-FLAG M2 Affinity gel (Sigma Aldrich). After 2 h of incubation, beads were washed 3 times with standard lysis buffer supplemented with extra NaCl (1 M final concentration). After washing, samples were split into two parts, and sample buffer with or without a reducing agent (β-mercaptoethanol) was added. Samples were boiled for 5 min and separated on a 15% polyacrylamide gel. For Western Blotting, proteins were transferred to immobilon-FL membranes. For diagonal electrophoresis, non-reducing samples were separated on a 15% polyacrylamide gel, after which the entire protein containing lane was excised, reduced, and separated on a new 15% polyacrylamide gel. For visualization of diagonal electrophoresis proteins were stained by SimplyBlue.

### Filter trap assay

HEK293T cells were transfected with FLAG-p16^INK4A^, FLAG-p16^INK4A^C72A or GFP as a control. After two days, cells were scraped in 100 mM N-ethyl-maleimide in PBS (37°C) to prevent post-lysis oxidation and collected by centrifugation at 1200 rpm for 3 minutes. Lysis buffer (50 mM Tris pH 7.5, 1% v/v Triton X-100, 1.5 mM MgCl_2_, 5 mM EDTA, 100 mM NaCl, Protease inhibitors (Aprotinin + Leupeptin), NaF, 100 mM Iodoacetamide) added for 10 minutes followed by centrifugation for 15 minutes at 14000 rpm in an Eppendorf microfuge. The pellet was resuspended in benzonase buffer (50 mM Tris-HCL pH 8.0, 1 mM MgCl_2_) and 125U of benzonase was added followed by incubation for 1hr at 37 degrees. The reaction was stopped by adding 2x termination buffer (40 mM EDTA, 0.2 % SDS, with or without 10 mM DTT). Protein content was measured using the BCA kit and 50 micrograms of protein per sample were loaded on a Bio Dot apparatus (Biorad) equipped with a 0.2 micron pore size nitrocellulose membrane soaked in buffer B (10 mM Tris-HCL pH 8.0, 150 mM NaCl, 2 % SDS) on top of two Whatmann paper filters soaked in Buffer A (10 mM Tris-HCL pH 8.0, 150 mM NaCl, 1 % SDS), after pre-washing the membrane with 100 µl of buffer A. Samples were pulled through the membrane using vacuum, followed by washing thrice with 100 µl of buffer A. The nitrocellulose membrane was further processed as for Western Blotting to detect trapped p16^INK4A^.

### Kinase assay

The kinase assays in this study were performed with immunoprecipitated FLAG-p16^INK4A^ or FLAG-p16^INK4A^C72A and the attached binding-partners. Beads were washed three times with standard lysis buffer followed by two washes with kinase buffer containing 25 mM Tris pH 7.5 and 10 mM MgCl_2_. The beads were subsequently incubated for 30 min at 30 °C in 20 µl kinase buffer supplemented with 1 mM DTT, 1 µg human GST-Rb (48-378 Sigma Aldrich) as substrate and 0.2 mM ATP. Directly after the assay beads were boiled for 5 min in reducing sample buffer and loaded on a 15% polyacrylamide gel followed by Western Blotting.

### Protein expression and purification

An *Escherichia coli* codon-optimized gene of *Homo sapiens* p16^INK4^ was cloned into a modified version of pETM-11 that includes a 6xHis, a protein A tag and a tobacco etch virus (TEV) protease cleavage site, leaving an additional glycine residue at the N-terminal cleavage site. For a glycerol stock solution, competent *E. coli* BL21 (DE3) cells were transformed with the p16^INK4^ harboring gene and inoculated in minimal medium in the presence of kanamycin (50 µg/l). After overnight incubation at 37 °C the cell suspension was mixed with glycerol to 50% (v/v) glycerol and stored at −80 °C. Uniformly ^15^N-labeled protein samples were prepared from this stock by growing the cells in minimal medium containing ^15^NH_4_Cl as the only nitrogen source. The protein synthesis was induced by addition of 0.5 mM isopropyl-1-thio-D-galactopyranoside (IPTG) at an OD_600_ of 0.8 and cells were harvested after 14 h of induction at 19 °C. Cells were resuspended in purification buffer (110 mM potassium acetate, 20 mM HEPES [4-(2-hydroxyethyl)-1-piperazineethanesulfonic acid], pH 8.0, 2 mM MgCl_2_, 2 mM β-mercaptoethanol (BME), 5% (v/v) glycerol and 20 mM imidazole) and ultrasonicated for lysis. The solution was applied to a gravity Ni-NTA agarose (QIAGEN) column following the manufacturer’s instructions. After washing with purification buffer including 20 mM imidazole, the protein was eluted by using purification buffer including 200 mM imidazole and further purified by gel chromatography on an ÄKTA pure system equipped with a HiLoad 16/600 Superdex 75 pg column (GE Healthcare) running with purification buffer. The fractions containing the protein were pooled, incubated with 0.2 mg TEV protease overnight at 4 °C while dialyzed against purification buffer using a dialysis membrane with 5 kDa molecular weight cut-off (ZelluTrans, Roth). On the following day, the protein solution was applied to the Ni-NTA agarose column again to remove the cleaved tag, remaining undigested protein and the His_6_-tagged TEV protease. The eluate was then buffer exchanged into purification buffer where BME was substituted with 1 mM DTT using dialysis. The protein solution was aliquoted, frozen in liquid nitrogen and stored at −80 °C. NMR spectroscopy showed that no differences were observed after the freezing and thawing of the samples.

### NMR spectroscopy

The protein stock was freshly buffer exchanged into NMR buffer (4 mM HEPES, 5 mM EDTA, pH 7.5) before measurement. Samples for protein characterization contained 1 mM DTT (dithiothreitol) whereas samples for redox-experiments did not. The protein concentration was 0.15 mM and contained 7% (vol/vol) ^2^H_2_O for the lock signal. ^1^H^15^N HSQC NMR spectra were measured on an Avance III 600 Bruker NMR spectrometer equipped with a cryogenic triple resonance probe with gradients in the z-direction. The recycle delay of the redox experiments performed (S-glutathionylation and hydrogen peroxide addition) was set to 1 s with a spectral window of 11/34 ppm in ^1^H/^15^N dimensions, 2048 points in the direct ^1^H dimension, 128 complex data points in the indirect ^15^N dimension, and with 8 or 16 scans per increment. Spectra were processed with the NMRPipe package ^55^ and analyzed by CcpNmr Analysis ^56^.

For determination of the S-glutathionylation redox potential, the concentration of GSSG in the sample buffer was gradually increased (0.2; 1.0; 2.3; 3.3; 3.9; 5.9; 7.8; 8.8; 9.8; 11.7 mM) by titrating from a 65 mM pH corrected stock solution while the concentration of the reduced form of GSH was kept constant at 4 mM. At each concentration point, a ^1^H^15^N HSQC NMR spectrum was recorded immediately after addition of the oxidation agent. For assignment of the S-GSHylated protein, HNCA and CBCACONH spectra were acquired. Determination of the redox potential was performed following the protocol of Piotukh *et al.* ^12^. In summary, peak intensities of well isolated peaks (N71, C72, D74, T79) of the oxidized form were normalized to a non-affected asparagine side chain resonance and plotted against the redox potential of the buffer. This potential was determined from the standard half-cell potential of the glutathione redox couple (−240 mV at pH 7.0, 25 °C) after pH corrections ^48^. The data were fitted to the sigmoidal decay function (equation 1):

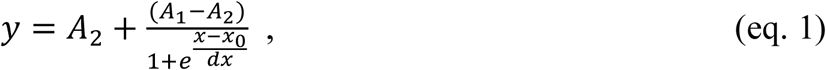

where A_1_ and A_2_ are the initial and final plateaus of the function, dx is the slope and x_0_ refers to the desired parameter of the redox potential.

For oxidized samples of p16^INK4^, a final concentration of 50 mM hydrogen peroxide (H_2_O_2_, from a 30% solution diluted with NMR buffer) or 100 µM diamide (from a freshly prepared 10 mM stock solution in H_2_O) was added to the ^15^N-labeled protein solution. ^1^H^15^N HSQC NMR spectra were acquired after the addition of the oxidizing agent in subsequent manner. After 10 h, no spectral changes were observed for WT protein anymore and the sample was applied to a size exclusion column (HiLoad 10/300 Superdex 75 pg column, GE Healthcare) and the dimeric fraction of about 50% of the total protein was pooled. The protein was applied to SDS gel electrophoresis in the presence and absence of BME.

### Small-angle X-ray scattering measurements

SAXS data was recorded on an in-house SAXS instrument (SAXSess mc2, Anton Paar, Graz, Austria) equipped with a Kratky camera, a sealed X-ray tube and a two-dimensional Princeton Instruments PI•SCX:4300 (Roper Scientific) CCD camera. Measurements were performed with 90 min exposure time (540 frames of 10 s each) of three concentrations of 2.5 mg/ml, 1.25 mg/ml and 0.625 mg/ml in NMR buffer. The sample of reduced p16^INK4A^ was measured immediately after purification and buffer exchange. The oxidized sample of p16^INK4A^ was treated overnight with 50 mM H_2_O_2_ at room temperature. Next day, the sample was applied to size exclusion chromatography and the peak from 9-10 ml elution volume was pooled (see Figure S6, B) and concentrated. Individual frames of the 90 min exposure were compared and no radiation damage was observed. A range of momentum transfer of 0.012 < s < 0.63 Å^-1^ was covered (s =4π sin(θ)/λ, where 2θ is the scattering angle and λ = 1.5 Å is the X-ray wavelength). SAXS data were analyzed with the package ATSAS version 2.8.1 ^57^. Desmearing of the oxidized sample could not be performed because the scattering curve showed strong aggregation.

### Thioflavin-T Fluorescence Assay

Samples of p16^INK4A^ (20 µM) were prepared in buffer (4 mM HEPES, pH 7.5) with 20 µM ThT. Three replicates were measured for each sample. Samples were subjected to 1 mm orbital shaking for 2 min of every 5 min, in a 96-well, half-area, fluorescence plate (Corning) and measurements were taken every 5 min. Samples were excited at 440 nm and fluorescence emission was measured at 480 nm. Measurements were made using a Tecan GENios microplate reader.

### Negative-stain Transmission Electron Microscopy

Protein samples were prepared by incubation of p16^INK4A^ samples with or without oxidizing agents for 6–24 h at RT. Copper grids with 300 meshes coated with formvar/carbon film (Electron Microscopy Sciences, Hatfield, USA) were glow-discharged in argon atmosphere for 30 s at 3 mA. Grids were floated on a 5 µl drop of a 50 µM protein sample and incubated for 60 s. Grids were washed once with water, and then floated on 5 µl of uranyl acetate solution (2% w/v) for 30 s. Micrographs were taken on either a JEOL JEM 100CX transmission electron microscope or a JEOL JEM 1400 Plus transmission electron microscope (JEOL, Tokyo, Japan).

### Prediction of aggregation-prone regions

Four different aggregation prediction programs, TANGO ^20^, PASTA ^21^, Zyggregator ^6^ and Aggrescan ^23^ were used to identify putative regions responsible for the beta-aggregation of p16^INK4A^ using the default settings.

### Congo-red absorbance

Absorbance spectra were measured on samples of p16^INK4A^ (20 µM) with Congo red (20 µM) in p16^INK4A^ buffer. Spectra were recorded from 400–650 nm, with a resolution of 2 nm, on a Tecan Infinite 200Pro microplate reader in a 96-well, half-area, fluorescence plate (Corning).

### Fourier-transform infrared spectroscopy

p16^INK4A^ (80 µM) samples were treated with 50 mM H_2_O_2_ overnight in p16^INK4A^ buffer, then dialysed against water overnight. Spectra were recorded on a JASCO FT/IR-4100 FT-IR spectrometer with attenuated total reflectance (ATR) attachment. The samples were measured with 128 scans at a resolution of 2 cm^-1^ at room temperature.

### Solid-state NMR spectroscopy

Approximately 10 mg (150 µM) of ^13^C-^15^N labelled p16^INK4A^ fibrils were prepared in p16^INK4A^ buffer by addition of 50 mM H_2_O_2_ and incubation at 37 °C overnight. The sample was packed into a 3.2 mm MAS rotor by ultracentrifugation. Spectra were recorded on a Bruker Avance III 750 MHz spectrometer (Bruker BioSpin) equipped with a 3.2 mm triple-resonance MAS probe. A proton-driven spin diffusion (PDSD) spectrum ^58^ was recorded at 16.5 kHz MAS, with 50 ms mixing time, at a set temperature of 273 K, with 352 scans and 22 ms and 8 ms acquisition time in the direct and indirect dimensions, respectively. Experiments were acquired using Topspin 3.2 (Bruker Biospin) and analysed using CCPN Analysis 2 ^56^ The secondary chemical shift was calculated as [δ(Cα_observed_) – δ(Cα_random coil_)] – [δ(Cβ_observed_) – δ (Cβ_random coil_)] with random coil chemical shifts taken from ^59^.

### Sample preparation for mass spectrometry

HEK293T cells were transfected with FLAG-p16^INK4A^ or FLAG-p16^INK4A^ C72A. After two days, half of the dishes (3 full 15cm dishes per sample) were treated for 10 minutes with diamide (250 µM) (Sigma Aldrich) and subsequently harvested for immunoprecipitation with FLAG beads as described before. After washing 2x with 1 M salt buffer and 3x with PBS to remove all soap, proteins were eluted from the beads by two times 5 minutes incubation with 75 µl 0.1 M glycine pH 2. The protein containing supernatant was transferred to a new tube and incubated for 20 minutes with 10 mM DTT and 2 M Urea (dissolved in 100 mM Tris pH 7.5), followed by 10 minutes incubation with iodoacetamide (50 mM). To digest the proteins, 0.25 µg trypsin (Promega) was added per sample and samples were incubated ON at 25 °C. The next day C18-stagetips were used for filtering and loading of the protein digest.

### Mass spectrometry and data analysis

The mass spectrometry was performed as previously described ^11^.

For the analysis Maxquant software version 1.5.1.0 ^60^ was used. During the analysis, oxidation of methionine, alkylation of cysteines with iodoacetamide were set as variable modification. Proteins were identified by the IPI human V3.68 database and the relative amounts of protein in the separate experiments, the Intensity Based Absolute Quantification (IBAQ) ^61^, as well as the label free quantification ^62^were calculated.

Further analysis was done using R version 3.4.0. Proteins identified with two or more unique peptides were filtered for reverse hits, decoy hits and standard contaminants and samples that show aberrant clustering or low protein count were excluded. Subsequently the IBAQ data was Log2 transformed and normalized for IP efficiency. For p-value calculations, left-censored missing data was first imputed using a stochastic minimal value approach. Imputation was performed by random draws from a Gaussian distribution centered in the 10^−4^ quantile of the known data with a standard deviation that is same as the observed values ^63^ A standard t-test was performed on the imputed data and p-values were adjusted for multiple testing using the Benjamini-Hochberg correction.

### Mass Spectrometry of GS-p16^INK4A^

The intact mass of p16^INK4A^ before and after oxidation by GSSG was obtained from LC-MS analysis, that was performed on recombinantly expressed protein (see protein expression and purification section). The sample was desalted using 10 kDa Amicon Ultra-0.5 centrifugal filter devices prior to measurement. 150 pmol of protein were injected into an Agilent 1200 system (Vienna, Austria) coupled to an LTQ-FT mass spectrometer (Thermo Fisher Scientific, Waltham, MA, USA). Separation was carried out on a monolithic PepSwift® column (500 µm x 5 cm Monolithic PS – DVB) at a flow rate of 20 µl/min using the following gradient, where solvent A is 0.05 % trifluoroacetic acid in water and solvent B is 0.05 % trifluoroacetic acid in acetonitrile: 0-5 min: 10 % B; 5-55 min: 10-100 % B; 55-65 min: 100 % B, 65-80 min: 10 % B. The sample was ionized in the electrospray source and analyzed using FT-MS operated in positive ion mode applying full scan MS (m/z 300 to 2000) with a resolution of 400,000. Acquired data were processed with the software Protein Deconvolution 2.0 SP2 (Thermo Fisher Scientific) using a S/N threshold of 5 and a relative abundance threshold of 20 %.

### Flow cytometry

U2OS cells were stably transfected with a Tet-repressor expressing construct as well as with pTON EGFP-p16^INK4A^ or pTON EGFP-p16^INK4A^C72A. To study cell cycle profiles by flow cytometry, A mixture of these cells and untransfected cells as controls was treated with doxycycline for 24 hrs to induce the expression of EGFP-p16^INK4A^ or EGFP-p16^INK4A^C72A. Cells were then treated overnight with nocodazole (0.25 µg/ml) (Sigma Aldrich) to trap all cycling cells in Mitosis (see scheme on Figure S1A) The next day cells were washed, trypsinized and fixed with cold ethanol (70%). After overnight incubation at 4°C, cells were centrifuged and resuspended in a mixture of propidium iodide (PI) (30 µM) (MPBio Life Sciences) and RNAse (200 µM/ml) (Sigma Aldrich) in PBS without CaCl_2_ and MgCl_2_ (Sigma Aldrich). Cells were gated for EGFP positivity and PI was measured in the FL3 channel and 10.000 EGFP expressing cells were gated per sample.

---

**Figure S1.**
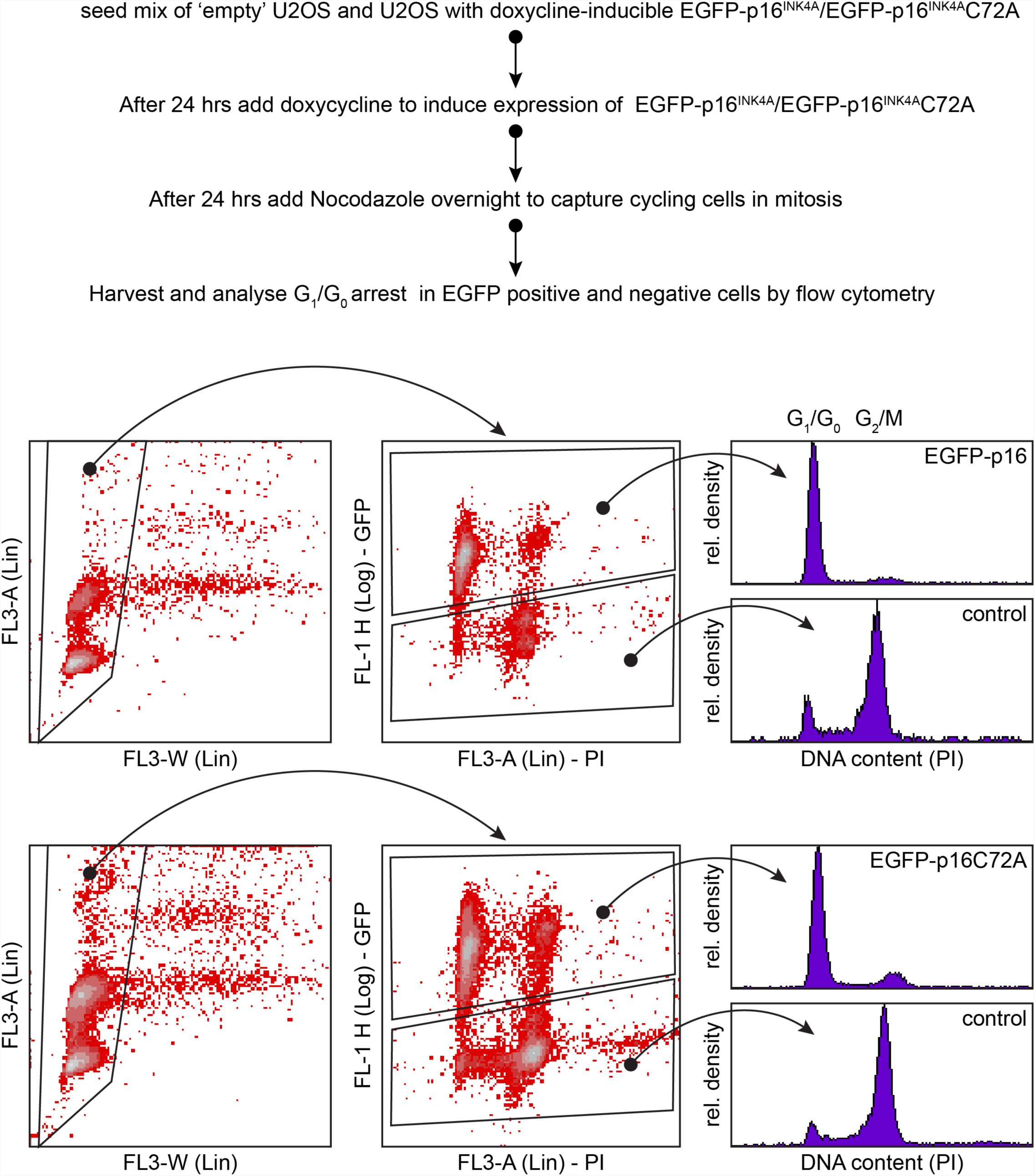
Cell cycle profiling of U2OS cells expressing doxycycline inducible EGFP-p16^INK4A^ wildtype or EGFP-p16^INK4A^C72A. U2OS without these constructs were mixed in the dish to serve as a control. Cells were trapped overnight in mitosis using Nocodazole to make G1/G0-arrested cells obvious from those that were still cycling. The C72A mutation leaves the cell-cycle inhibitor function of p16^INK4A^ intact.

**Figure S2.**
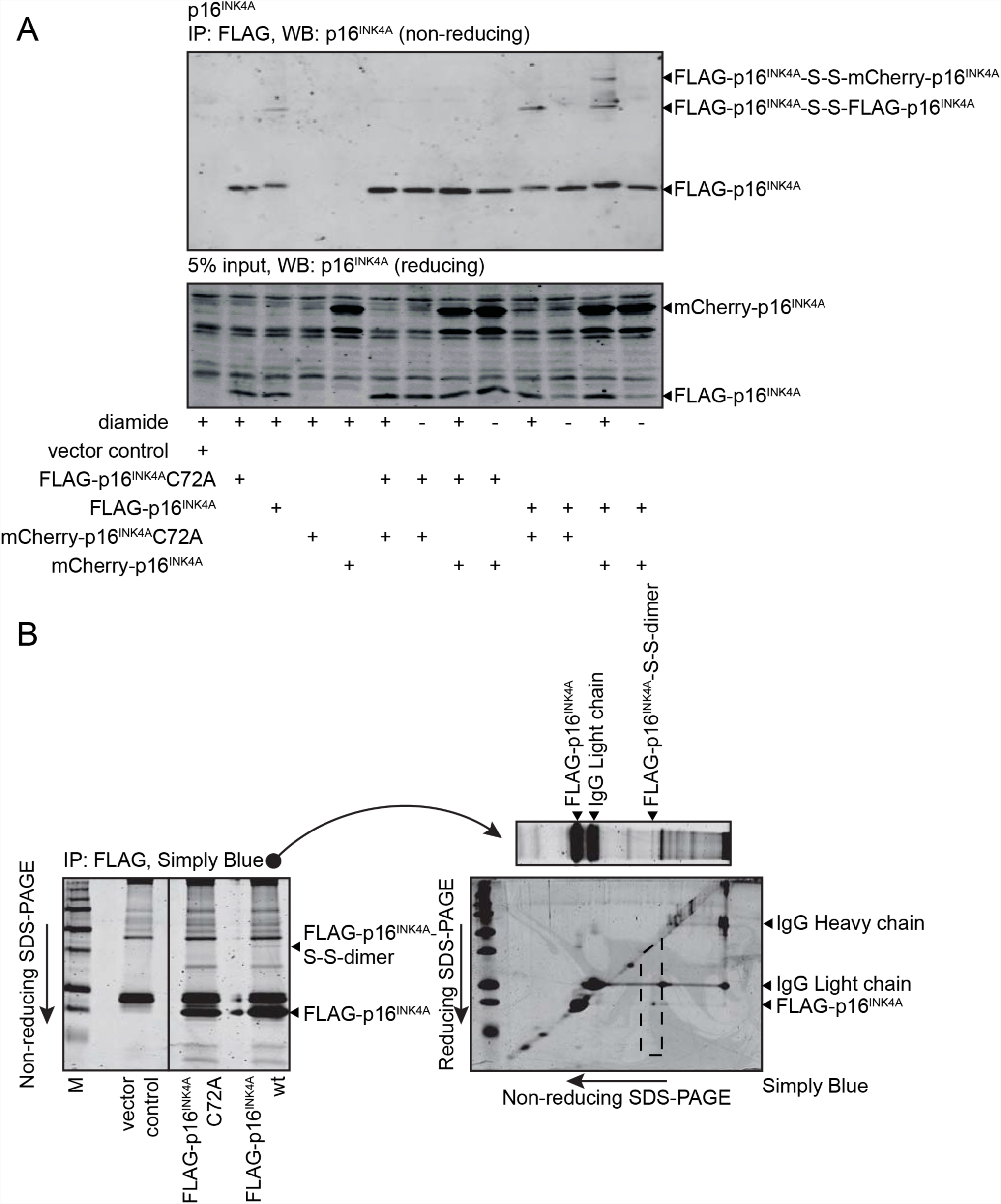
SDS-PAGE analysis of p16^INK4A^. A) mCherry-p16^INK4A^ co-immunoprecipitates with Flag-p16^INK4A^ in a disulfide-dependent manner. Dimers consisting of both two Flag-p16^INK4A^ molecules or one Flag-p16^INK4A^ and one mCherry-p16^INK4A^ molecule can be detected. B) Diagonal electrophoresis (non-reducing SDS-PAGE followed by reducing SDS-PAGE) shows that the high-molecular weight form of Flag-p16^INK4A^ separates in a single dot under the diagonal, affirming that the disulfide-containing high-molecular weight species of p16^INK4A^ consists of only p16^INK4A^ oligomers. Collectively, and because p16^INK4A^ has only one cysteine, these data must mean that the slow-migrating species of p16^INK4A^ detected under non-reducing conditions are indeed p16^INK4A^ homo-dimers.

**Figure S3.**
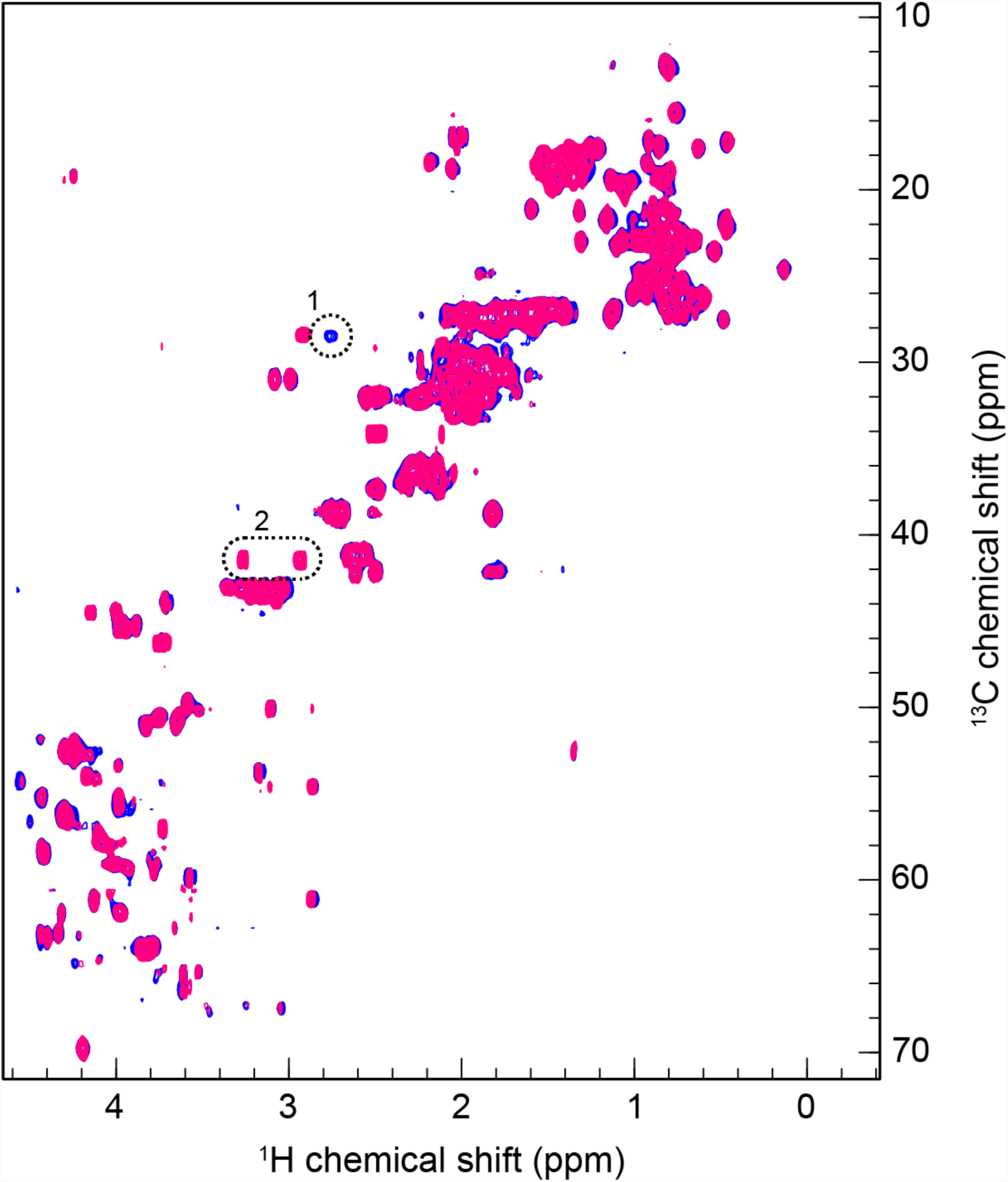
Overlap of ^1^H^13^C HSQC spectra in the reduced (blue) and glutathionylated state (magenta) The blue peak labeled as (1) is the reduced C72 Cβ resonance. After titration with glutathione to the end point, the resonance disappears and the oxidized Cβ resonance appears in (2).

**Figure S4.**
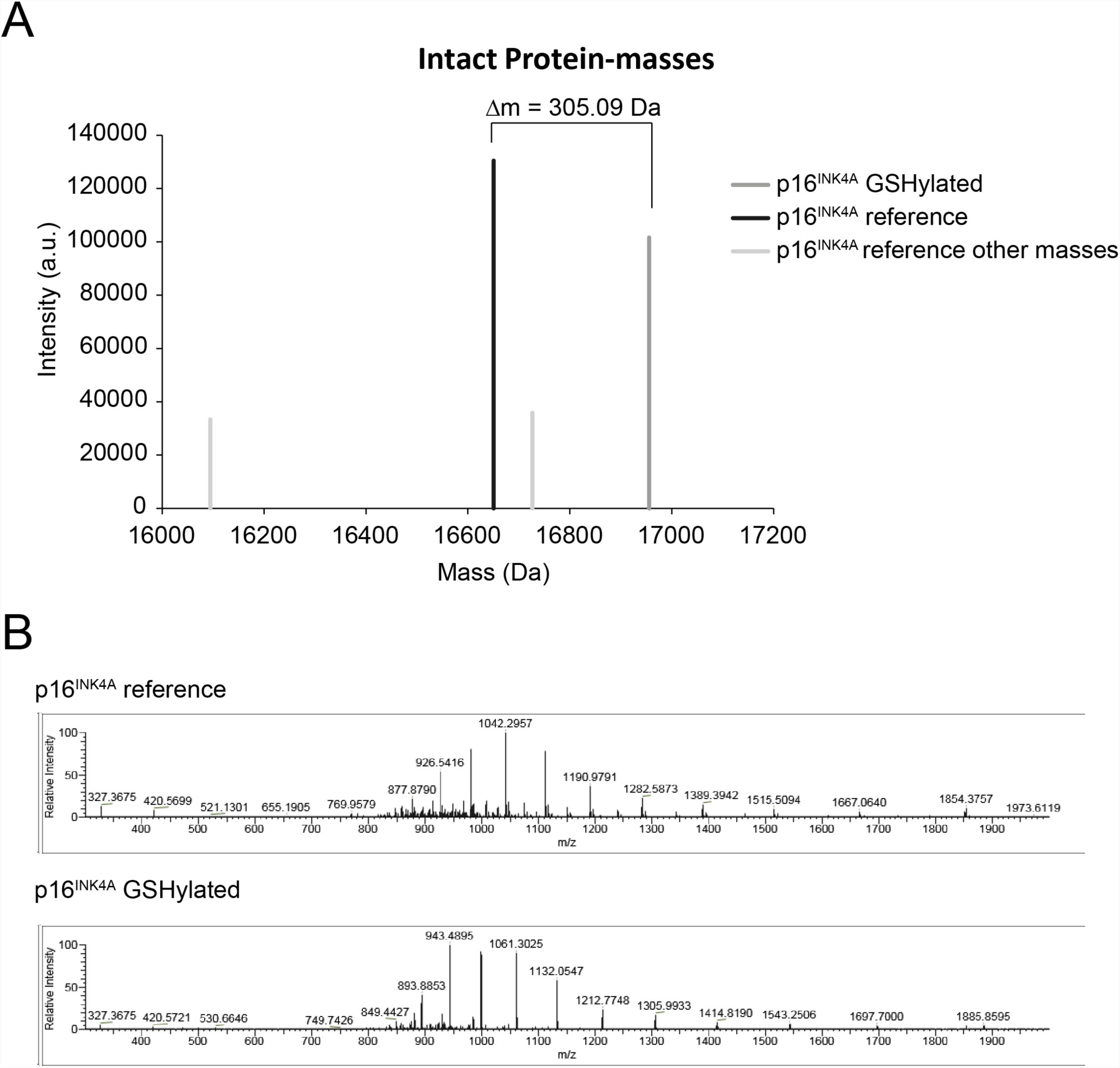
Mass spectrometry. Mass spectrometry analysis of reduced and glutathionylated p16INK4A. (A) Deconvoluted mass spectra of p16INK4A treated with GSSG shows a mass-shift of 305 Da which is in agreement with the addition of disulfide-linked Glutathione to p16INK4 C72. (B) m/z spectra of the LC-MS analysis of recombinant p16INK4A before and after GSHylation.

**Figure S5.**
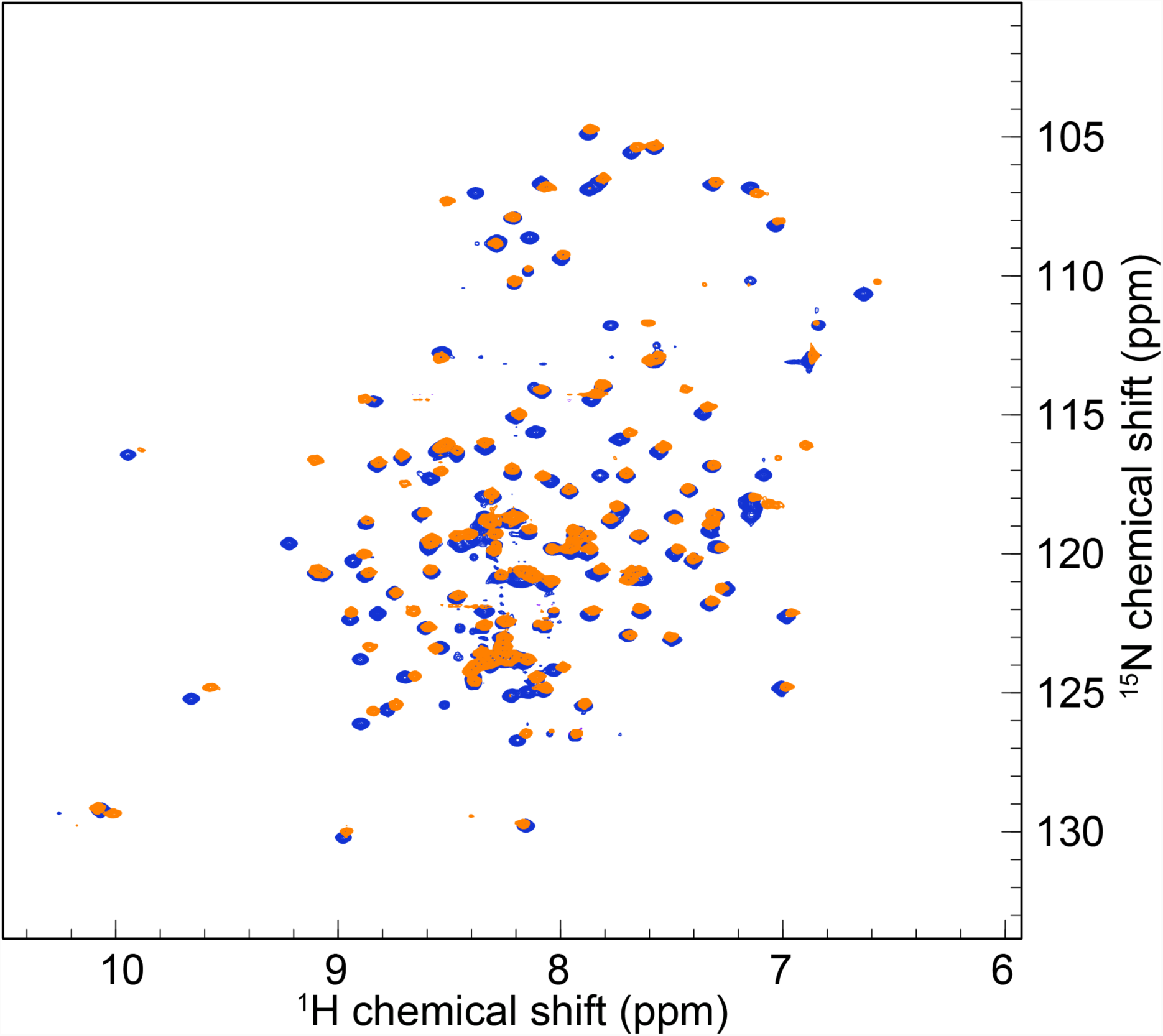
^1^H^15^N HSQC solution NMR spectra of WT p16^INK4A^ (blue) and p16^INK4A^ C72S (orange).

**Figure S6.**
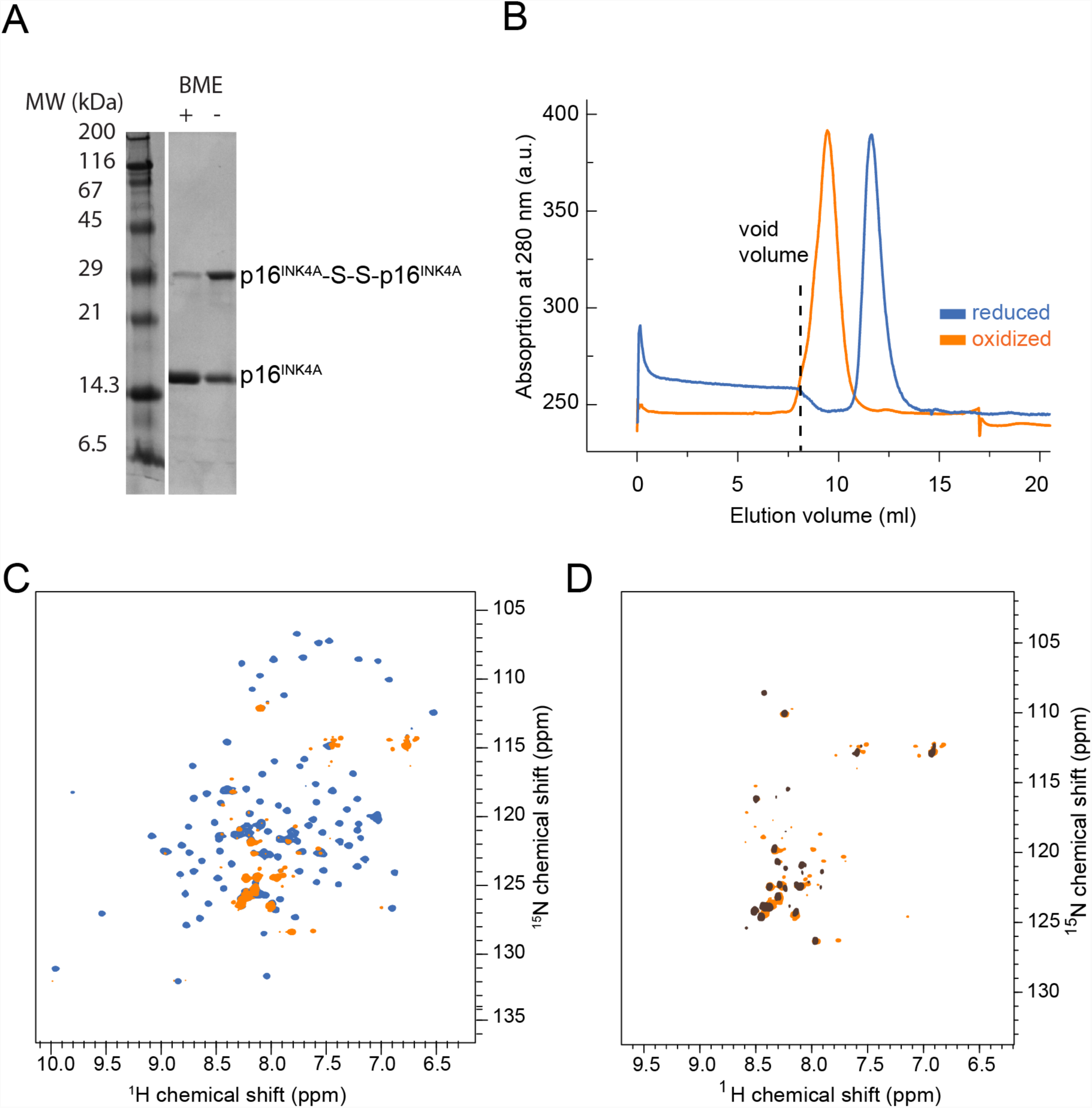
Analysis of oxidized p16^INK4A^. (A) SDS-PAGE of wild-type p16^INK4A^ after hydrogen peroxide oxidation in the presence (left) and absence of reducing agent BME in the sample buffer. (B) Size exclusion chromatography of WT p16^INK4A^ before (blue) and after (orange) overnight oxidation with 50 mM hydrogen peroxide at room temperature. (C) Overlap of ^1^H^15^N HSQC spectra of reduced WT p16^INK4A^ and after addition of diamide (orange). (D) Overlap of ^1^H^15^N HSQC spectra of WT p16^INK4A^ oxidized with H_2_O_2_ (brown) and diamide (orange).

**Figure S7.**
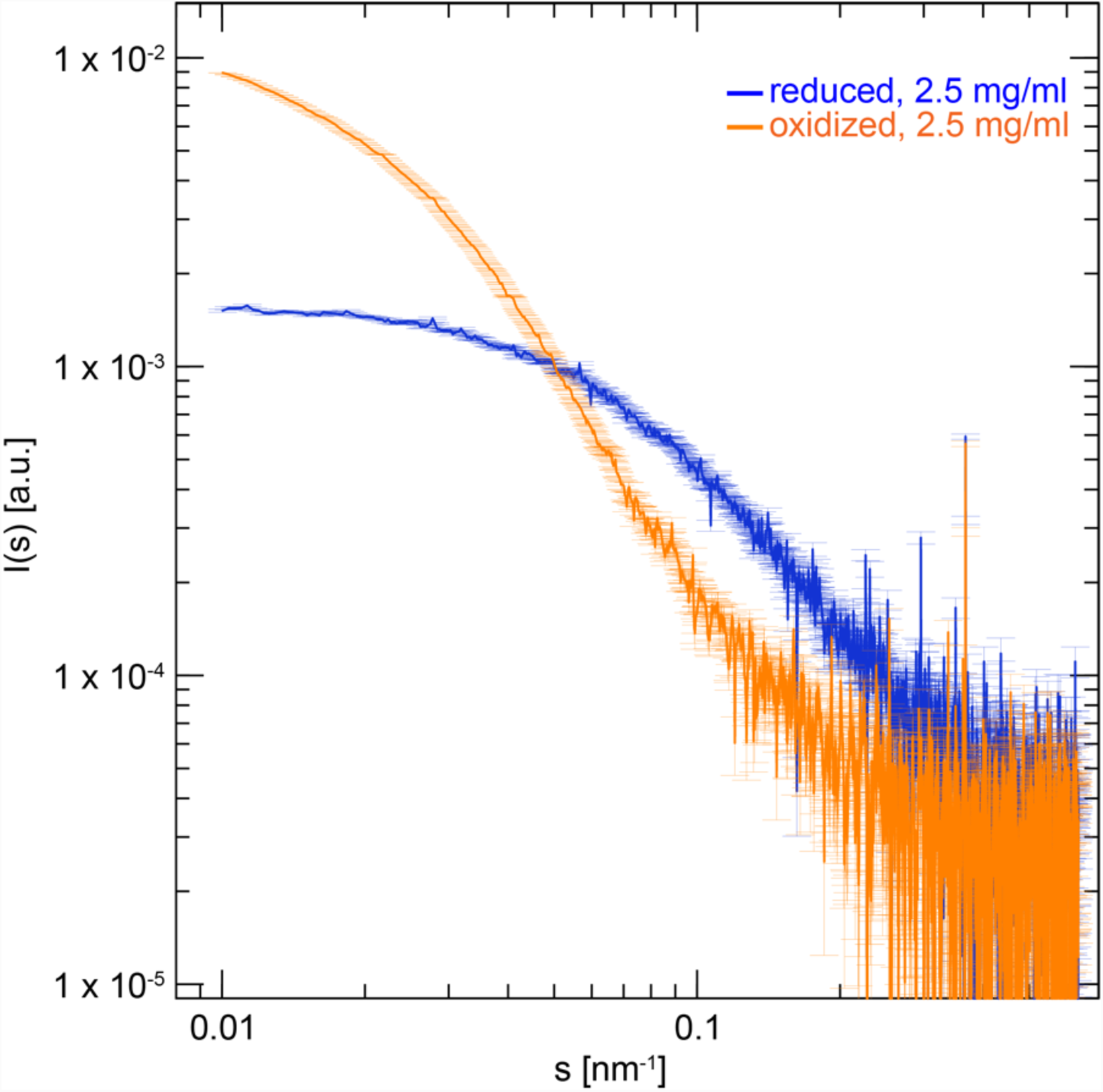
SAXS curves of reduced (blue) and oxidized (orange) p16^INK4A^ samples. Scattering curves of 2.5 mg/ml protein concentration are presented. The oxidized sample displays typical characteristics of an aggregated protein state and could not be desmeared for detailed data analysis.

**Figure S8.**
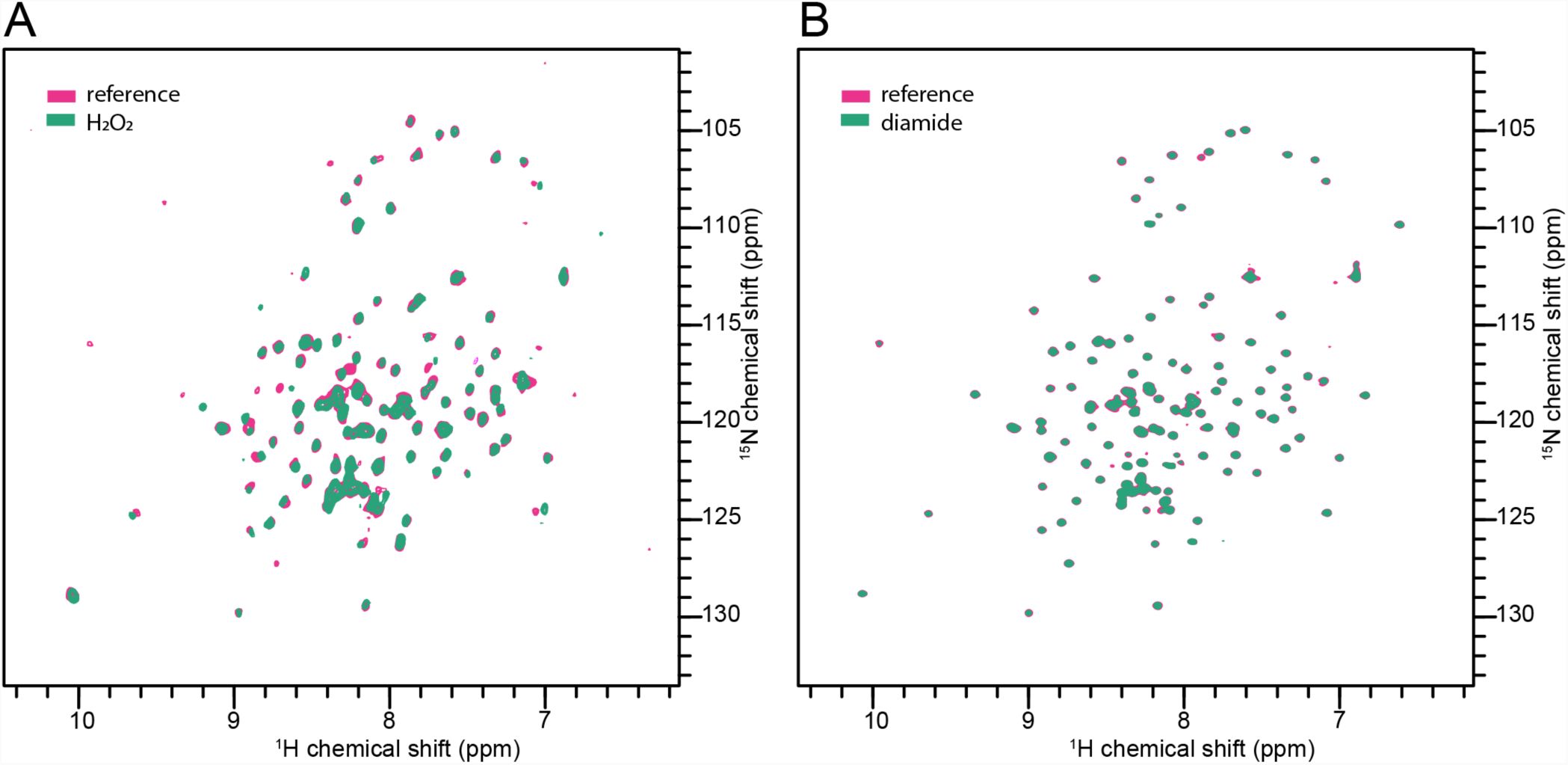
Effect of oxidizing agents of GS-p16^INK4A^. (A) Overlap of ^1^H^15^N HSQC spectra of reduced (magenta) and H_2_O_2_ treated GS-p16^INK4A^ (B) Overlap of ^1^H^15^N HSQC spectra of reduced (magenta) and diamide treated GS-p16^INK4A^

**Figure S9.**
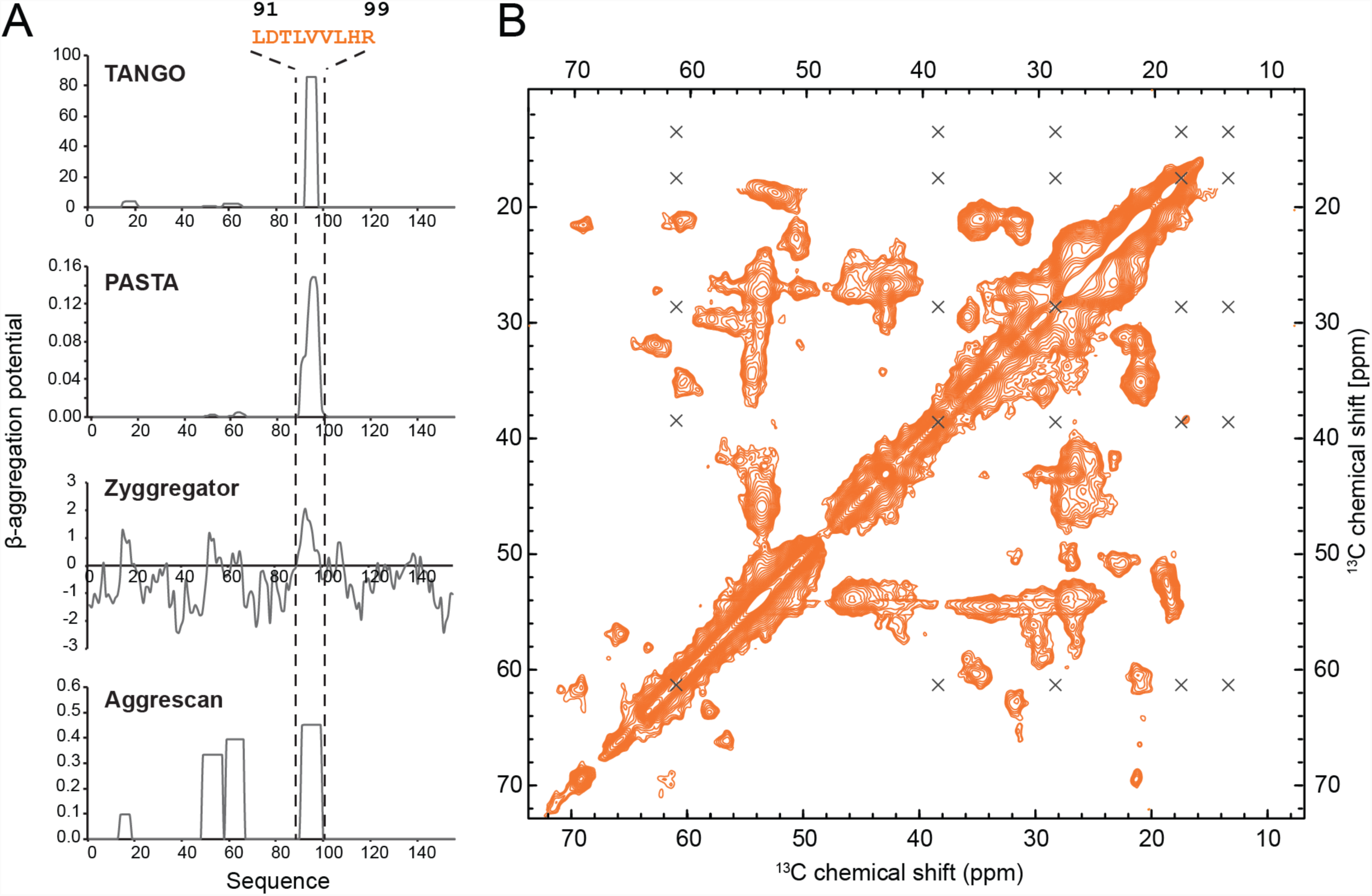
Predictions of p16^INK4A^ aggregation regions. (A) Four beta-aggregation prediction algorithms predict the p16^INK4A^ sequence from about residues 90 to 99 have high aggregation propensity. (B) ^13^C^13^C solid state NMR spectrum of p16^INK4A^ fibrils showing full aliphatic region. Crosses mark positions where peaks would be expected arising from isoleucine residues. There are three isoleucines in the full p16^INK4A^ sequence, so these residues are therefore absent from the folded beta-sheet core of p16^INK4A^ fibrils.

**Figure S10.**
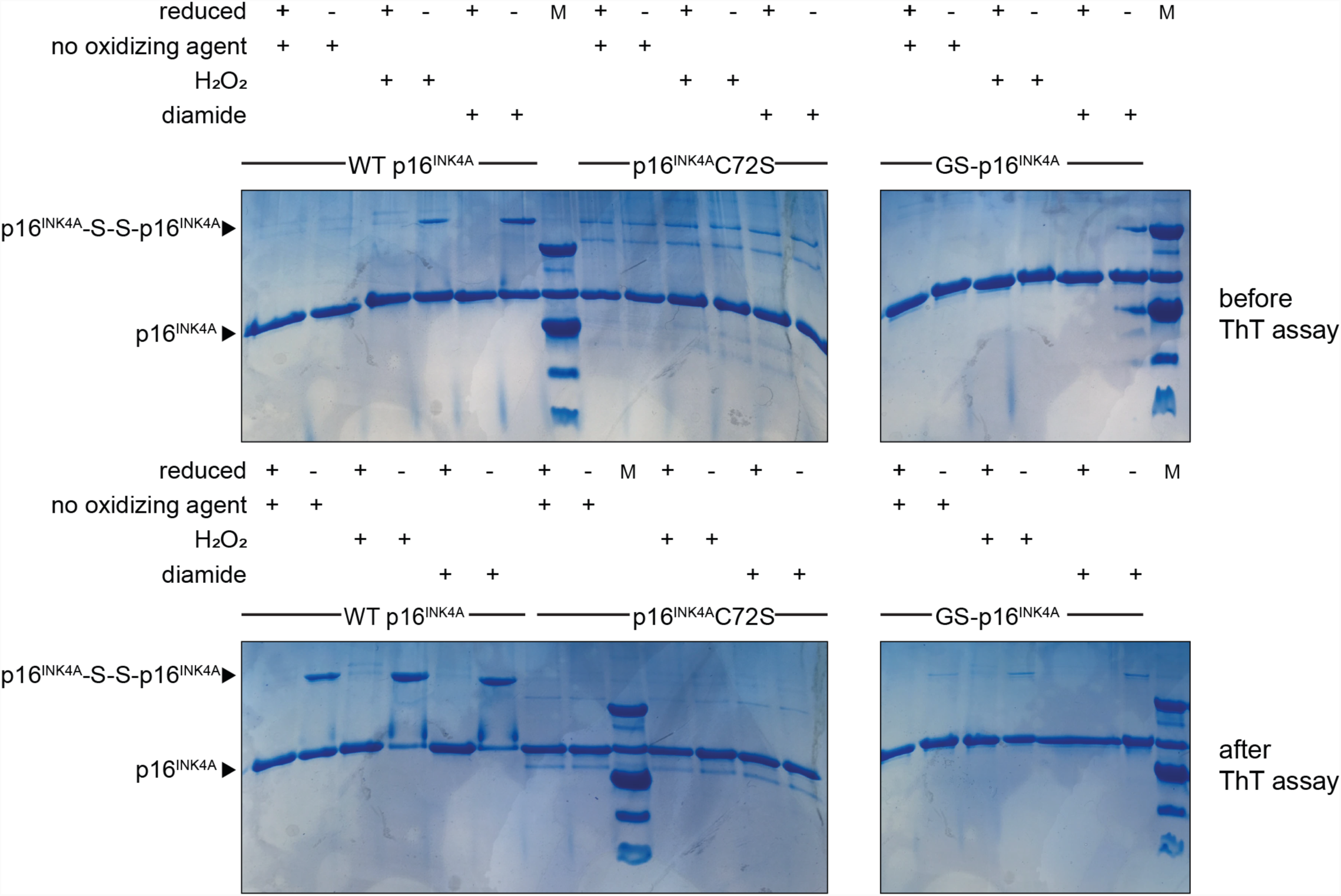
SDS-PAGE of p16^INK4A^ samples before and after fibril formation. Samples were taken before and after a ThT kinetics assay (see Figure 5B) and analysed by SDS-PAGE. WT p16^INK4A^, p16^INK4A^C72S and GS-p16^INK4A^ were allowed to form fibrils in the absence or presence of oxidizing agents (H_2_O_2_ or diamide). Samples for gel analysis were incubated is SDS loading buffer with or without reducing agent (DTT). Dimer bands can be seen in non-reduced samples of WT p16^INK4A^, especially after oxidizing agent treatment and after incubation during ThT assay.

